# Ancient balancing selection maintains incompatible versions of a conserved metabolic pathway in yeast

**DOI:** 10.1101/829325

**Authors:** James Boocock, Meru J Sadhu, Joshua S Bloom, Leonid Kruglyak

## Abstract

Differences in nutrient availability have led to the evolution of diverse metabolic strategies across species, but within species these strategies are expected to be similar. Here, we discovered that the galactose metabolic pathway in the yeast *Saccharomyces cerevisiae* exists in two functionally distinct, incompatible states maintained by ancient balancing selection. We identified a genetic interaction for growth in galactose among the metabolic genes *GAL2, GAL1/10/7*, and *PGM1*. We engineered strains with all allelic combinations at these loci and showed that the reference allele of *PGM1* is incompatible with the alternative alleles of the other genes. We observed a strong signature of ancient balancing selection at all three loci and found that the alternative alleles diverged from the reference alleles before the birth of the *Saccharomyces sensu stricto* species cluster 10-20 million years ago. Strains with the alternative alleles are found primarily in galactose-rich dairy environments, and they grow faster in galactose, but slower in glucose, revealing a tradeoff on which balancing selection may have acted.

## Introduction

To grow and reproduce, organisms must extract chemicals from their environments and convert them into energy and cellular building blocks. Variation in nutrient availability between environments has led to the evolution of diverse and complex interconnecting metabolic enzymatic pathways. In humans, mutations in these pathways give rise to diseases known as inborn errors of metabolism (*1*). The budding yeast *Saccharyomyces cerevisiae* has been extensively used as a model system for unraveling the genetic and biochemical basis of eukaryotic metabolism (*2*). Although *S. cerevisiae* prefers glucose as a carbon source, it can utilize a wide variety of other sugars, and work over many decades has identified and characterized the regulatory and enzymatic components of pathways that process various carbon sources (*3*).

One exceptionally well-studied example of alternative carbon utilization is the Leloir pathway of galactose metabolism (*4*). This pathway consists of a galactose transporter, encoded by the gene *GAL2*, enzymes that work together to convert galactose into glucose-1-phosphate, and regulatory components that control their expression. The enzymatic components are encoded by *GAL1, GAL10*, and *GAL7*, together known as the structural galactose genes. Phosphoglucomutase, encoded by *PGM1* and *PGM2*, then converts glucose-1-phosphate to glucose-6-phosphate—the substrate for glycolysis. The *GAL* genes are strongly repressed when either glucose or glycerol is present but are induced up to 1000-fold when only galactose is available (*5*). Induction of the galactose pathway is controlled by the *GAL4* transcription factor and its regulators *GAL80* and *GAL3* (*4*).

Although the enzymes involved in galactose metabolism are highly conserved from yeast to humans, the regulation of galactose metabolism varies substantially (*6, 7*). Within the yeast family *Saccharomycetaceae*, species show radically different responses to galactose (*8*). For example, in *S. uvarum*, a species that is separated from *S. cerevisiae* by approximately 10-20 million years, *GAL* genes are not repressed by glucose (*9, 10*). Some strains of *S. kudriavzevii* can metabolize galactose, while others have lost this ability through pseudogenization of multiple genes in the pathway, and it has been proposed that the two versions of the pathway have been maintained by multi-locus balancing selection (*11*). Recent population surveys of *S. cerevisae* identified substantial sequence diversity in genes involved in galactose metabolism and uncovered some strains that lack glucose repression, a feature thought to be characteristic of galactose regulation in this species (*12–15*).

In the course of mapping the genetic basis of variation in growth on multiple carbon sources, we discovered a three-way genetic interaction for growth in galactose (*16*). Here, we fine-mapped these three loci to genes in the galactose pathway and identified specific allelic combinations of these genes that are incompatible for growth in galactose. We characterized the global distribution of these alleles in over 1000 yeast strains (*12, 13*) and found that most strains contain one of two functionally distinct versions of the galactose metabolic pathway that have been maintained by ancient balancing selection. We experimentally identified a fitness trade-off between these versions which provides a possible explanation for the maintenance of these alleles.

## Results

### Genetic mapping identifies a three-way genetic interaction for galactose growth

We previously used a panel of ∼14,000 haploid *S. cerevisiae* progeny, derived from crosses of 16 parental strains, to map over 4,500 quantitative trait loci (QTLs) that influence growth in 38 different conditions (*16*). When we searched for non-additive QTL interactions, one higher-order interaction stood out by virtue of its large effect size and high statistical significance. This interaction involved three loci and influenced growth in galactose for progeny from a cross between a soil isolate, CBS2888, and a clinical isolate, YJM981 (three-way effect size 0.19, P<10^−15^; Figure 1a; Table 1; Supplementary Table 1). The same interaction was replicated in the second cross that had CBS2888 as a parent (three-way effect size 0.19, P<10^−12^), but it was not observed in any of the other 14 crosses, suggesting that it was driven by CBS2888 alleles (Supplementary Figure 1). The three QTLs are located on different yeast chromosomes (II, XI, and XII) and, as expected, segregate independently (chi-square test, P=0.20). Together, these QTLs account for 57% of the phenotypic variance of growth in galactose, with 36% of the variance explained by additive effects and 21% by interaction effects. The non-additive nature of the effects of the three loci is best illustrated by the phenotype of segregants that inherit the CBS2888 allele at the loci on ChrII and ChrXII and the non-CBS2888 allele at the locus on ChrXI— these segregants grow much more slowly in galactose than do those with any other combination of alleles (Figure 1a).

**Table 1.**
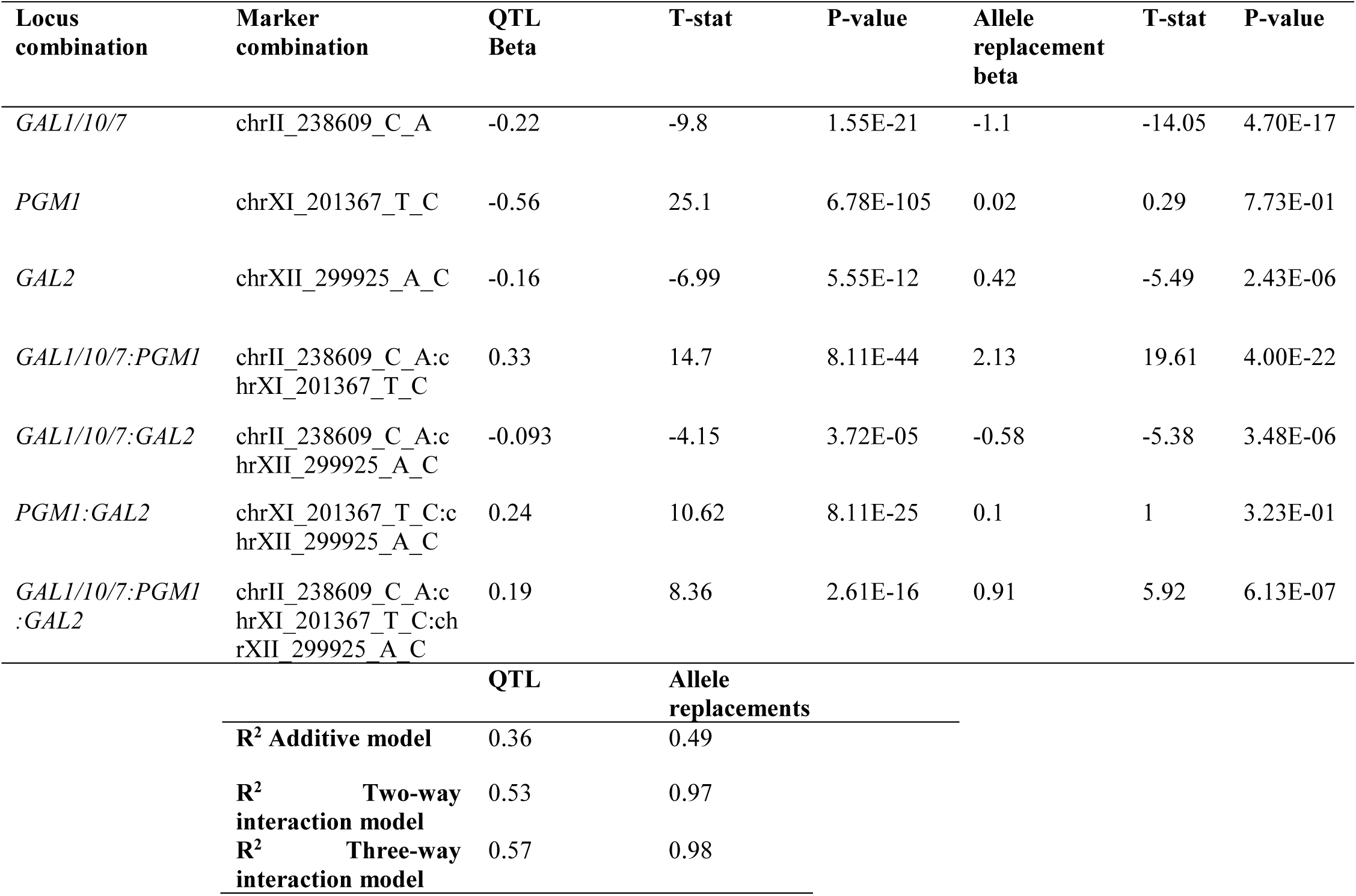
Additive, two-way, and three-way linear model coefficients and fits for segregants and allele replacement strains when grown in galactose.

**Figure 1.**
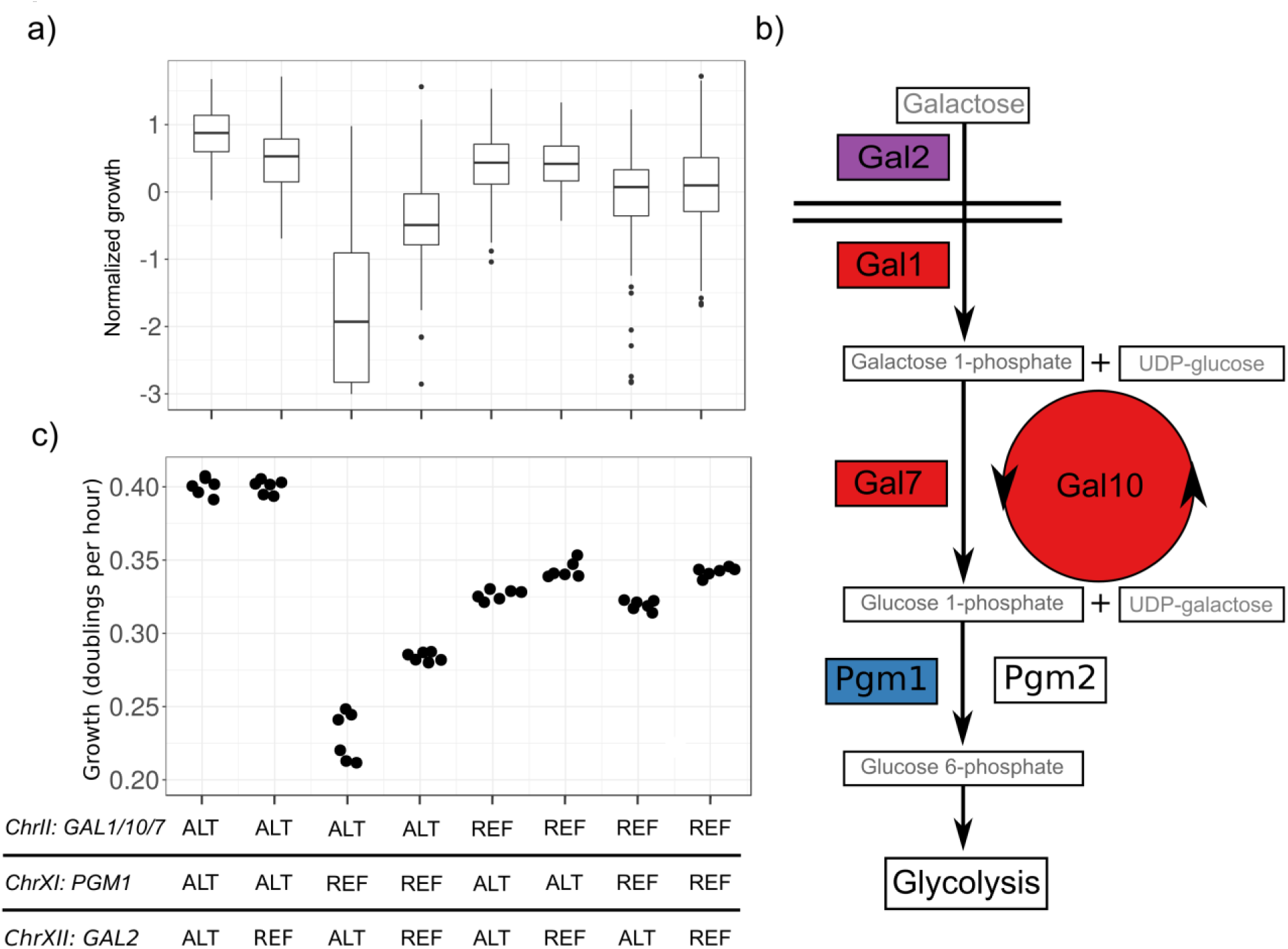
Three-locus genetic interaction for growth in galactose. a) Boxplots show growth of yeast segregants on agar plates containing 2% galactose as the sole carbon source. Each boxplot corresponds to segregants with one of eight distinct combinations of alleles at the three loci (ChrII: *GAL1/10/7*, ChrXI: *PGM1*, and ChrXII: *GAL2*). b) The galactose metabolic pathway. Components of the pathway corresponding to the three loci are shown in different colors. c) Growth of allele replacement strains in galactose. Each dot shows a biological replicate. The strain genotypes correspond to the segregant groups in a. BY alleles are designated as REF; CBS2888 alleles are designated as ALT.

### The causal genes are members of the galactose metabolic pathway

We examined the three QTL confidence intervals in detail and observed that all three contained genes that encode components of galactose metabolism (Figure 1b) (*4*). The ChrII QTL contained *GAL1, GAL10*, and *GAL7*, the ChrXI QTL contained *PGM1*, and the ChrXII QTL contained *GAL2*. We also found that the CBS2888 alleles of these genes were highly diverged from those of the other 15 parental strains in our previous study (*16*), with less than 80% sequence identity with the reference genome for coding and promoter regions of *GAL1, GAL2, GAL7*, and *GAL10*, and 94% sequence identity for the coding region but only 48% for the promoter region of *PGM1*. In contrast, coding regions and promoters of genes from the CBS2888 genome have an average sequence identity with the reference genome of 99.3% and 98.6%, respectively (Supplementary Figure 2). The CBS2888 allele of the *GAL2* locus contains a duplication of *GAL2* (Supplementary Figure 3). We hereafter refer to the divergent galactose alleles found in CBS2888 as the alternative alleles, and the alleles observed in the other strains as the reference alleles.

**Figure 2.**
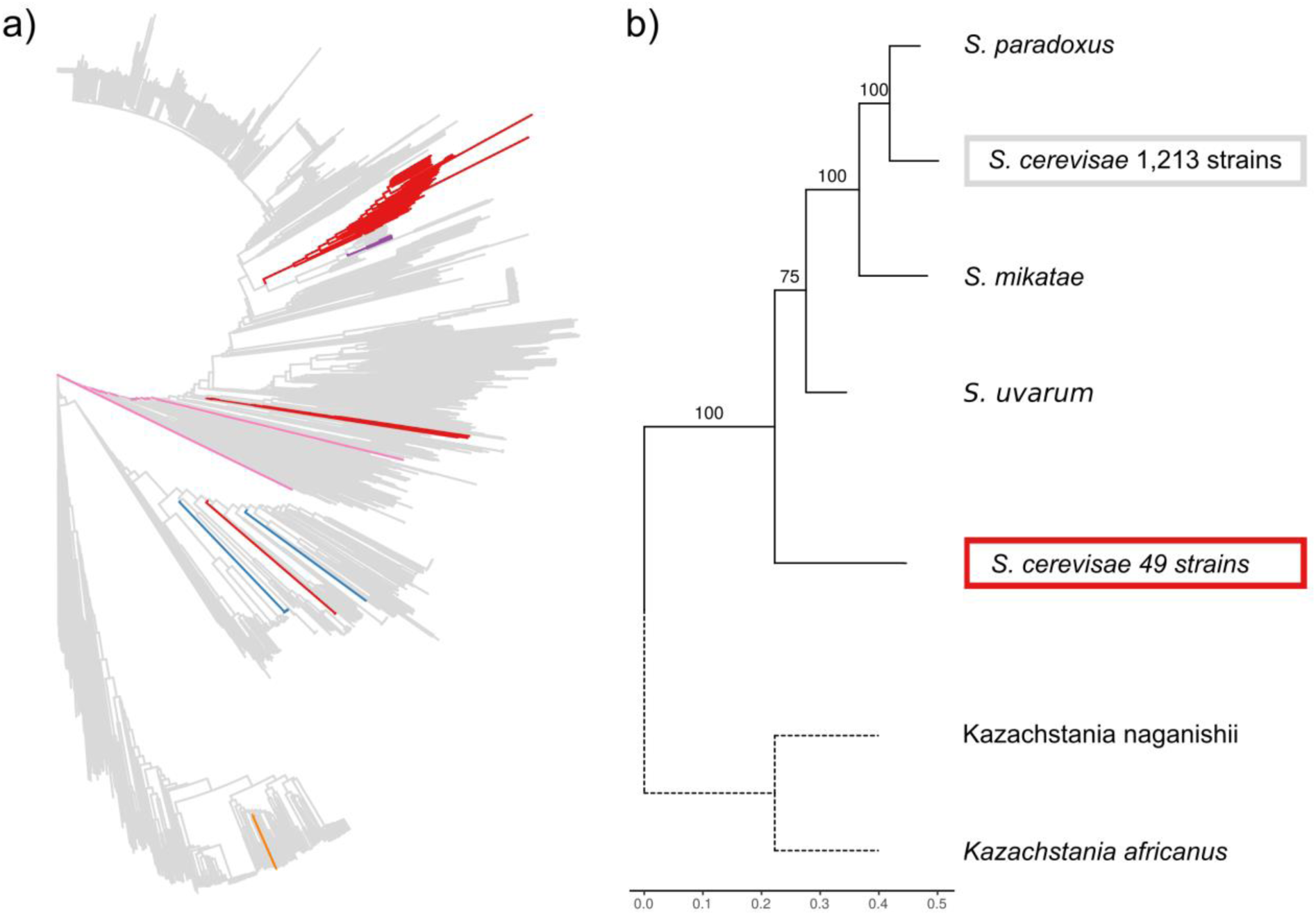
The alternative galactose alleles are broadly distributed and fall outside the *Saccharomyces sensu stricto* species complex. a) Genome-wide neighbor-joining tree of 1,276 sequenced yeast isolates. Clusters of branches with the same genotypes at the three galactose loci are colored as follows: all three reference alleles (grey), all three alternative alleles (red), alternative *GAL1/10/7* allele and Chinese *PGM1* and *GAL2* alleles (blue), reference alleles of all genes except *GAL7* (purple), heterozygous at the *PGM1* promoter and homozygous for the reference alleles at the other loci (pink), and a deletion of the entire region containing *GAL1/10/7* and reference alleles at the other loci (orange). b) Phylogenetic tree of the *GAL1/10/7* alleles from CBS2888 (alternative), BY (reference), other members of the *Saccharyomyces sensu stricto* and two outgroup species. Bootstrap support in percent is shown for each clade. Scale bar shows the estimated number of substitutions per site. The outgroup branches (dotted lines) were rescaled to the average branch length.

**Figure 3.**
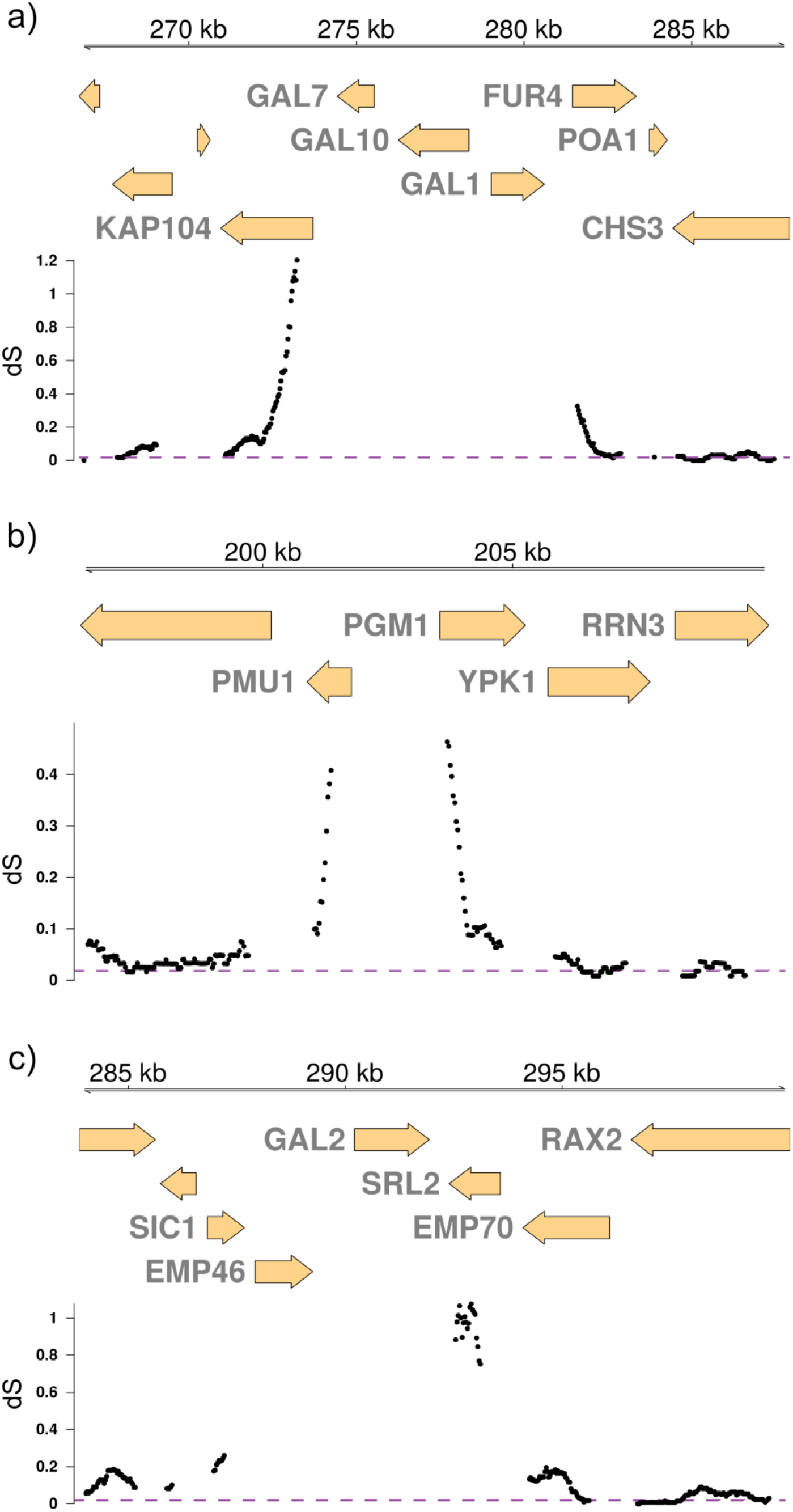
A signature of ancient balancing selection. Estimated rate of synonymous substitutions per site (dS) between CBS2888 (alternative) and BY (reference) is shown for genes surrounding the galactose loci. Estimates of dS are plotted as dots for 200-codon windows stepped every ten codons. a) genes adjacent to *GAL1/10/7*, b) genes adjacent to the *PGM1* promoter, c) genes adjacent to *GAL2*. dS was not estimated for *EMP46* due to the presence of an early stop codon interrupting this gene in CBS2888. The purple dashed line shows the genome-wide average dS of 0.014.

To test whether variants in these genes underlie the QTL signals, we used CRISPR-Cas9 to engineer strains with all eight possible combinations of the three alternative and three reference galactose alleles in a common genetic background (Methods; Supplementary Tables 2-4)(*17*). For the *GAL1/10/7* and *GAL2* loci, we replaced these genes and their intergenic regions in a laboratory strain with the alternative regions from CBS2888, including the *GAL2* duplication (Supplementary Figure 3). For the *PGM1* locus, we replaced only the promoter region based on the observed divergence pattern described above. We measured the growth rates of the eight engineered strains in galactose and found that the results recapitulated those from QTL mapping (Figure 1b; Supplementary Figure 4; Table 1). The strain with all three alternative alleles attained the highest growth rate in galactose (0.40 doublings per hour), growing 17% faster than the strain with all reference alleles (0.34 doublings per hour. P<10^−7^). The strain with the combination of the reference *PGM1* promoter allele and the alternative alleles of the other *GAL* genes exhibited a severe growth defect and had the lowest growth rate of all the engineered strains (0.23 doublings per hour, 43% slower than the strain with all three alternative alleles, P<10^−6^). These results confirm that variants in the coding and intergenic regions of *GAL1/10/7* and *GAL2* and in the promoter region of *PGM1* are responsible for the observed QTL interaction. Further, we find that the reference *PGM1* promoter allele is incompatible with the alternative *GAL1/10/7* and *GAL2* alleles.

**Table 2.** Summary statistics for alignments between the alternative and reference galactose alleles.

| Gene/region | % identity | dS | 95% C.I |
| --- | --- | --- | --- |
| *GAL1* | 76.1 | 2.12 | 1.52-3.92 |
| *GAL10* | 76.7 | 2.61 | 1.66-3.79 |
| *GAL7* | 77 | 2.59 | 1.50-3.61 |
| *GAL2* | 77.9 | 2.38 | 1.56-4.15 |
| *PGM1* promoter | 46.7 | NA | NA |

**Figure 4.**
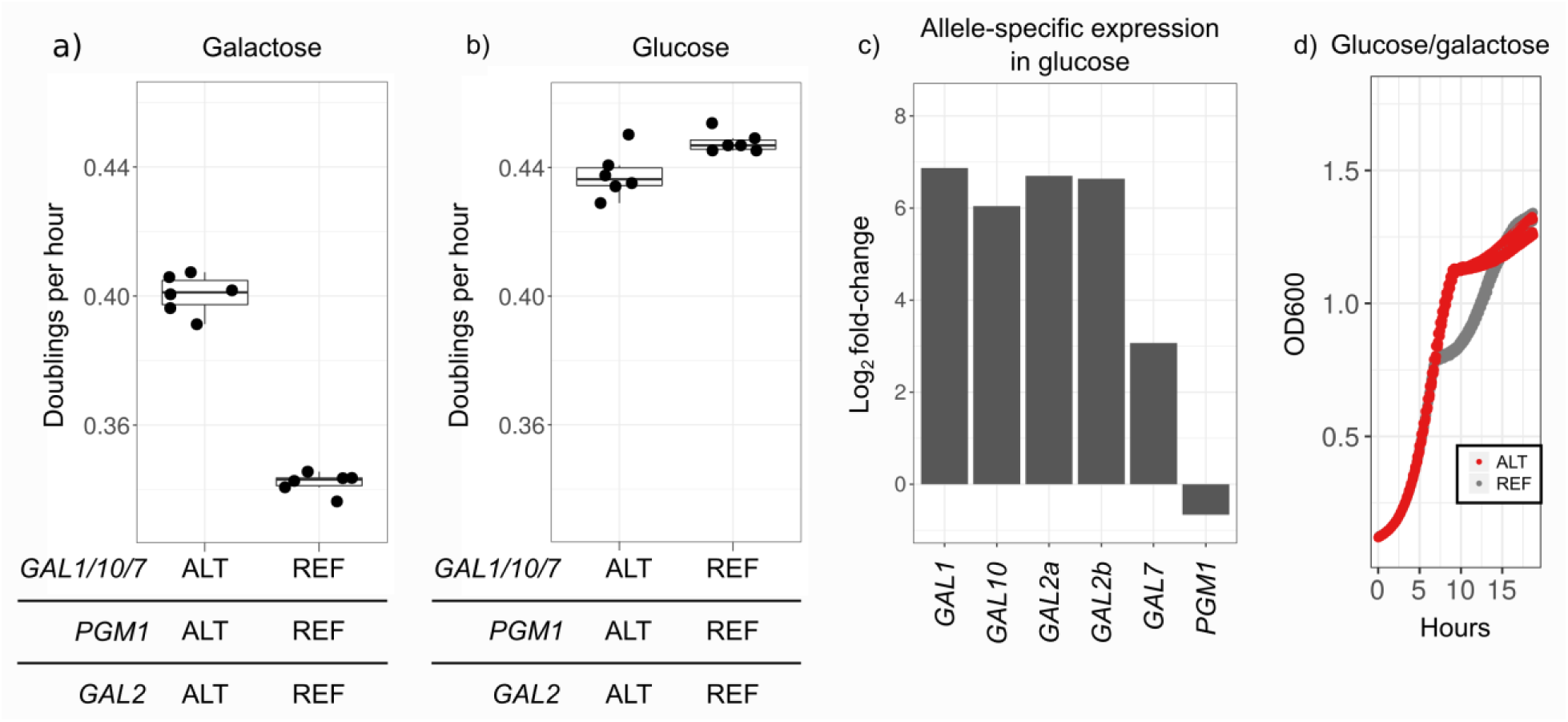
Trade-offs between the alternative and reference alleles in the galactose pathway. a) Growth rates of allele replacement strains with all three reference (right) or all three alternative (left) alleles in galactose as a sole carbon source. Biological replicates are shown as dots and summarized in boxplots. b) As in a, but for cells grown in glucose. c) Allele-specific expression of a diploid hybrid (CBS2888xBY) strain grown in glucose. The bar plot displays log_2_ fold-changes for the alternative alleles of the galactose genes and *PGM1*, as compared to the reference genes. The centromere-proximal copy of *GAL2* in CBS2888 is denoted as *GAL2a*, and the other copy as *GAL2b.* d) Growth curves of allele replacement strains with all three alternative (red) or reference (grey) galactose alleles in mixed glucose/galactose medium demonstrate the presence of the diauxic shift in strains with the reference but not the alternative pathway.

### The alternative galactose pathway requires galactose-responsive *PGM1* expression

To better understand *cis*-acting regulatory differences between the alternative and reference galactose alleles (*18*), we grew a diploid hybrid strain (CBS2888xBY) in glucose, transferred it to galactose medium, and sequenced RNA from samples collected throughout a growth time course. In glucose medium, the expression of the CBS2888 allele of *PGM1* was slightly lower than that of the reference allele. In contrast, one hour after the switch to galactose, the expression of the CBS2888 allele of *PGM1* was 15.5-fold higher than that of the reference allele (P<10^−100^), and this difference persisted for the rest of the time course (Supplementary Figure 5, Supplementary Table 5ab). The alternative *PGM1* promoter allele contains a *GAL4* upstream activating sequence (UAS)(*19*), whereas the reference allele does not (Supplementary Figure 3). We engineered a point mutation disrupting the UAS in a strain with all three alternative galactose alleles. This single mutation recapitulated the severe growth defect in galactose observed in a strain with a combination of the reference allele of the PGM1 promoter and alternative alleles of the other *GAL* genes (Supplementary Figure 6). We conclude that induction of *PGM1* in galactose, mediated through a *GAL4* UAS, is critical for the proper functioning of the alternative galactose pathway.

### The incompatible combination of galactose alleles is not found in nature

We were surprised to find two functional, but incompatible, versions of a conserved metabolic pathway co-existing in a single species. To learn whether strains with the incompatible combination of galactose alleles are found in nature, we searched for the alternative and reference galactose alleles in two large collections of sequenced *S. cerevisiae* isolates from around the globe, together comprising 1,276 strains (Figure 2a)(*12, 13*). We identified three most common combinations of galactose alleles: only reference alleles (1213 strains), only alternative alleles (49 strains), and 8 strains from China with the alternative *GAL1/10/7* allele and alleles of *GAL2* and the *PGM1* promoter that differ from both the reference and the alternative alleles (Supplementary Table 6). In addition, one strain had a deletion of the entire region containing *GAL1/10/7*, three strains contained reference alleles at all loci except *GAL7*, and two strains were heterozygous at the *PGM1* promoter and homozygous for the reference alleles at the other loci, indicating that strains with alternative and reference galactose alleles have not been completely reproductively isolated from each other. This is also demonstrated by the phylogenetic distribution of the isolates with alternative *GAL* alleles (Figure 2a). We found no strains with the reference *PGM1* promoter allele and the alternative *GAL1/10/7* and *GAL2* alleles—the combination which causes a severe growth defect in the lab—suggesting that this combination causes a fitness disadvantage in natural environments and has been purged by selection. This hypothesis is further supported by a high linkage disequilibrium index (*ϵ*=0.59) for the three loci (Supplementary Figure 7)(*20*). In comparison, random trios of single-nucleotide polymorphisms with a similar frequency (4-5%) have an average *ϵ* of 0.041 (95% C.I.=0.008-0.18).

We examined the sources of isolation of the 1,276 strains and found that the alternative galactose alleles are fixed in two lineages of strains found in domesticated dairy environments, including Camembert cheese from France, kefir grains from Japan, and fermented yak and goat milk from China (Figure 2a; Supplementary Table 6). These environments are rich in lactose, a disaccharide of glucose and galactose. *S. cerevisiae* does not metabolize lactose directly, but rather relies on the activity of other fungi and bacteria to break it down into glucose and galactose (*14*). Our observation that the alternative alleles improve growth rate in galactose suggests that these alleles are maintained by natural selection in dairy environments.

One hypothesis for the origin of the alternative galactose alleles in *S. cerevisiae* is a recent introgression around the time of domestication of milk-producing animals (*15*). We searched the genomes of all sequenced yeast species and did not find any that could have donated the alleles. We also noted that the alternative galactose alleles are found in strains that were not isolated from domesticated dairy environments, including CBS2888, which is a South African soil isolate, as well as human clinical isolates from Europe and soil and forest isolates from China. The global distribution of these alleles outside of dairy lineages and the lack of an obvious donor species suggested that they may have a more ancient origin. Maintenance of the alternative alleles in some non-dairy environments can be explained by galactose availability varying substantially among such environments (*21, 22*).

### Ancient balancing selection has maintained the galactose alleles of *S. cerevisae*

One force that can maintain highly diverged alleles within a species is balancing selection. This process is expected to generate a signature of elevated sequence divergence at linked neutral sites that decays as a function of genetic distance from the selected variant (*23*). We examined the rate of synonymous substitutions per site (dS) across the CBS2888 genome relative to the reference and observed a strong signature of ancient balancing selection at all three galactose loci (Figure 3a-c; Supplementary Notes 1 and 2). No comparable signatures were seen at other genomic loci in the CBS2888 genome (Supplementary Figure 8).

To date the divergence between the alternative and reference galactose alleles, we combined the estimated number of synonymous substitutions per site (dS) with the measured mutation rate µ=3.8×10^−10^ per site per generation in yeast (*24*). Under neutral theory, the dS value of 2.4 for the galactose alleles corresponds to a split between these alleles approximately 3.2 billion generations ago (95% C.I.=2.5-4.5 billion generations), which pre-dates the most recent common ancestor of the *Saccharomyces sensu stricto* species complex (*25*). This date is over 100 times older than the divergence between CBS2888 and the reference strain based on the average number of synonymous substitutions per site (0.014) for all genes (Supplementary Figures 9-10, Table 2, Supplementary Table 7). We note that three other genes in CBS2888 (*NTC20, BSC4, RFX1*) are estimated to have dS>1.0, but lack an associated signature of balancing selection at nearby genes. An ancient origin of the alternative galactose alleles is further supported by phylogenetic clustering analysis, which placed the alternative galactose alleles outside the *Saccharomyces sensu stricto* species complex (Figure 2b; Supplementary Notes 3 and 4)(*26*).

### The alternative and reference galactose alleles are advantageous in different environments

Balancing selection can act on fitness trade-offs, in which alleles with higher fitness in one environment have lower fitness in another(*23*). Although all strains grow faster in glucose than in galactose, strains with all three alternative galactose alleles grow faster in galactose than do strains with reference alleles (Figure 1c; Figure 4a), providing a fitness advantage on which selection could act in galactose-rich environments. Yeast encounter and metabolize a wide variety of sugars (*27*), but they prefer glucose (*28*). In glucose, strains with all three reference alleles grow 2% faster than strains with all three alternative alleles (T=-3.12, P=0.017) (Figure 4b, Supplementary Figure 11). This faster growth provides an explanation for maintenance of the reference alleles in strains that do not frequently encounter galactose.

We next looked for a mechanism that could underpin the observed growth differences. In strains with reference alleles, the *GAL* genes are robustly repressed by glucose and induced by galactose (*4*). The repression in glucose leads to a pause in growth known as the diauxic shift after the yeast consume all the glucose and must switch to metabolizing galactose. We found that strains with the three alternative galactose alleles do not undergo a diauxic shift during the switch from glucose to galactose (Figure 4d, Supplementary Figure 12ab). To investigate whether this difference is due to *cis*-regulatory changes, we examined RNA sequencing data collected for the diploid hybrid strain (CBS2888xBY) over the course of a transfer from glucose to galactose. In glucose, the reference alleles are repressed as expected, whereas the alternative *GAL* alleles are constitutively expressed (fold-change=40.6, P<10^−16^; Figure 4c; Supplementary Figure 13; Supplementary Table 5b). Constitutive expression of the *GAL* genes eliminates the diauxic shift (*9*), providing a fitness benefit when galactose is frequently encountered. However, gene expression can be costly (*29*), and constitutive expression of the alternative version of the galactose pathway may explain why it leads to a growth disadvantage in glucose.

## Discussion

We discovered that the key components of the galactose pathway in the yeast *Saccharomyces cerevisiae* (*GAL1/10/7, GAL2*, and *PGM1*) exist in two distinct functional but incompatible allelic versions. The alternative alleles of these genes are highly diverged from the reference alleles, and they are found primarily in isolates from galactose-rich environments, such as cheese, kefir, and milk. We show that balancing selection has maintained the two versions of the pathway for millions of years. We discovered a trade-off between the two versions—strains with the alternative alleles grow faster in galactose but slower in glucose. Carbon source composition varies substantially between natural environments (*21, 22*), and we propose that the availability of galactose determines the relative fitness of strains carrying the alternative alleles, providing a mechanism on which balancing selection has acted.

The alternative alleles of *GAL1/10/7* and *GAL2* that we identified show high sequence similarity to alleles previously found (*12–14*) and recently characterized in Chinese isolates (*15*). It was proposed that that *S. cerevisiae* acquired the alternative *GAL1/10/7* and *GAL2* galactose alleles through introgression approximately 10,000 years ago, around the time humans domesticated milk-producing animals, but no species that could have donated these alleles was identified (*15*). The divergence pattern we observe near the alternative alleles is not consistent with a relatively recent introgression, and instead supports the conclusion that the alternative and reference galactose pathways have been maintained by multi-locus balancing selection for millions of years. Balancing selection has typically been observed at single loci. Multi-locus balancing selection is expected to be rare because it has to overcome independent segregation of alleles at the different loci, therefore requiring strong selective forces, and few empirical examples have been reported previously (*11*).

By carrying out QTL mapping and searching for non-additive interactions, we were able to identify a third key component of the alternative version of the galactose pathway—the galactose-responsive *PGM1* promoter. The yeast family *Saccharomycetaceae*, which contains *S. cerevisiae*, has experienced recurrent gains and losses of galactose-responsive phosphoglucomutase (Pgm) genes(*8*). Most yeast species that lack a galactose-responsive Pgm grow slower in galactose, and the addition of a *GAL4* UAS in the promoters of *PGM1/2* in such species improves galactose growth(*8*). Species with a galactose-responsive Pgm generally grow faster in galactose, and the removal of the *GAL4*-UAS in these species decreases growth. Kuang et al. propose that galactose-responsive Pgm in these species improves growth in galactose by matching increased flux through the upstream components of the pathway (*8*). Our results suggest that another reason for repeated evolution of galactose responsive Pgm is that a metabolic bottleneck at this point in the pathway causes toxicity that leads to severe fitness defects.

In *S. uvarum*, a species that is separated from *S. cerevisiae* by approximately 10-20 million years, *GAL* genes are not repressed by glucose (*9, 10*). Some of the differences in glucose repression between *S. uvarum* and *S. cerevisiae* are attributable to the promoters of the structural galactose genes and *PGM1*. The alleles of the alternative galactose pathway of *S. cerevisiae* share similarities with the galactose pathway of *S. uvarum*. In particular, both pathways are not repressed by glucose, have a *GAL4-UAS* in the *PGM1* promoter, and have maintained a duplicated *GAL2*. These observations suggest that the alternative galactose pathway originated in a common ancestor of *S. cerevisiae* and *S. uvarum*, illustrating that the within-species differences we found in *S. cerevisiae* can provide insights into the evolution of this pathway across yeast species.

There are notable similarities between the incompatible allele combinations we identified and yeast models for human disorders of galactose metabolism. One out of every ∼45,000 children is born with classical galactosemia, an inborn error of metabolism caused by recessive mutations in *GALT*, the human homolog of *GAL7* (*30*). Individuals with galactosemia can develop life-threatening symptoms if galactose is not eliminated from their diet and often develop serious long-term complications, even with treatment. Classical galactosemia is known to result in the accumulation of galactose and certain metabolites, including galactose-1-phosphate, and the depletion of others, such as UDP-galactose and UDP-glucose (*31*). These metabolic changes, especially the accumulation of galactose-1-phosphate, are thought to be responsible for the severe pathologies observed in galactosemia, but the precise molecular mechanisms are not well understood (*31*). In yeast models of galactose toxicity, overexpression of the galactose enzymes (*GAL1/10/7*) results in accumulation of galactose-1-phosphate and reduced growth in galactose (*32*), as does inhibition of phosphoglucomutase with lithium (*33, 34*). We observe a similar reduction in growth in strains that combine an allele of *PGM1* that lacks galactose induction with galactose-adapted alleles of *GAL1/10/*7, which suggests that this incompatibility arises from the same metabolic defect that underlies galactosemia. The naturally occurring variants of the galactose pathway in *S. cerevisiae* may provide an avenue for uncovering the molecular mechanisms of this inborn error of metabolism in humans.

## Methods

Unless otherwise specified, all computational analyses were performed in R (v3.6.1)(*37*).

### Quantitative trait mapping

We obtained the additive QTL for each of the 38 phenotypes that were measured in ∼14,000 progeny from 16 parental crosses in Bloom et al (*16*). We tested all triplets of these QTL to see whether they were involved in any three-way QTL interactions. Specifically, for unique triplet combination of QTLs for each trait and cross we built a linear model with all additive QTLs, two-way QTLs, and the three-way QTL interaction terms. We tested whether this model fit significantly better than a nested model that included only additive QTLs and two-way interaction terms using a likelihood ratio test. We considered any three-way interaction with a q-value less 0.1 to be significant. The coefficient of variation, *R*^2^, was used to quantify how much variance in the phenotype was explained by the different QTL models. Boxplots of the normalized growth rate were made using ggplot2 (v3.2.0)(*38*). A chi-square test was used to evaluate whether the tree galactose loci segregated independently.

### Yeast strains

Strains, plasmids, and primers used in this study are listed in Supplementary Tables 2-4. To generate allele replacement strains, we used a two-guide RNA CRISPR system to introduce double-strand breaks flanking our regions of interest and provided linear repair templates of the desired replacement allele. The precise details for the construction of each of the strains are described below.

We engineered a lab yeast strain derived from BY4741 (YLK3221: Mata met15Δ his3Δ1 leu2Δ0 ura3Δ0 nej1Δ::KanMX) to generate strains with all eight combinations of the CBS2888 and BY4741 galactose alleles (*17*). To engineer strains with the CBS2888 *PGM1* promoter we used a plasmid that contained galactose inducible *CAS9* (PLK77, p415-GalL-Cas9-CYC1t)(*39*). For the CBS2888 *GAL2* and *GAL1/10/7* allele, we used a plasmid that contained a constitutively expressed *CAS9* (PLK91, pRS414-TEF1p-Cas9-CYC1t-NatMx). For the *PGM1* locus, we replaced the BY4741 *PGM1* promoter sequence with the CBS2888 *PGM1* promoter allele. We generated a repair template that contained the 2.9kb CBS2888 promoter sequence flanked by homology arms that were identical to the ends of the nearest flanking genes, *PMU1* and *PGM1*. For the *GAL2* locus, we replaced the BY4741 *GAL2* sequence and the surrounding non-coding region with the CBS2888 *GAL2* allele. We generated a repair template that contained the 5.7kb CBS28888 GAL2 sequence flanked by homology arms that were identical to the ends of the nearest flanking genes *EMP46* and *SRL2*. For the *GAL1/10/7* locus, we replaced the BY4741 *GAL1/10/7* sequence and the surrounding non-coding region with the CBS2888 *GAL1/10/7* allele. We generated a repair template that contained the 7.3kb CBS2888 *GAL1/10/7* sequence flanked by homology arms that were identical to the ends of the nearest flanking genes *KAP104* and. We co-transformed these repair templates with selectable plasmids expressing two guide-RNAs that exclusively cut near the 3’ and 5’ ends of the BY4741 region. We picked colonies and confirmed they had the exact intended replacement allele sequences using multiple sanger sequencing reactions (YLK3267, YLK3268, YLK3269).

We mated strains with the CBS2888 *GAL2* and *PGM1* promoter to obtain a heterozygous diploid (YLK3270), which we sporulated to get a haploid strain with both CBS2888 alleles. This strain was then mated to a strain with the CBS2888 GAL1/10/7 allele to obtain a heterozygous diploid (YLK3271). We also took care to ensure that auxotrophies and the drug marker were homozygous in this strain. We sporulated this strain and PCR genotyped the progeny. From this cross, we obtained two isolates of each of the eight possible combinations of the CBS2888 and BY4741 galactose alleles.

We made a point mutation in the GAL4-binding site of the CBS2888 promoter in strains with the CBS2888 *GAL1/10/7, PGM1* promoter, and *GAL2*. We used a plasmid that expressed a single guide RNA, which was co-transformed with a repair template. We confirmed that strains had the desired mutation using sanger sequencing (YLK3288-3291).

### Growth measurements

All growth experiments were performed at 30 degrees in YP media (2% bacto-peptone, 1% yeast extract) supplemented with 2% glucose, 2% galactose, or 1% glucose/1% galactose. Strains were always incubated with fast shaking in a BioTek Synergy™ 2 plate reader. Before each experiment, strains were grown to saturation in our plate reader in 96-well plates (Corning, Flat Bottom with Lid, #3370) in 2 % glucose. Strains were then diluted 1:100 into new 96-well plates and transferred to our plate reader, which automatically took optical density measurements (OD600) measurements every 15 minutes.

### Growth rate calculations

Growth rate was quantified as the geometric mean rate of growth (GMR). Our procedure for calculating the GMR follows that described in Brem et al (*9*). Briefly, a spline was fit in R using the splinefun function, and the time spent (*t*) between OD 0.2 and 0.8 was calculated. The GMR was then estimated as the log(0.8/0.2) /*t*. We then converted this GMR into doublings per hour. In each experiment, we measured the growth of three biological replicates of each strain. To determine whether the diauxic shift differed between our allele replacement strains, we grew these strains in 1% glucose/1% galactose medium and calculated the GMR between 0.8 and 1.1. This range of OD captures the range over which the reference strain pauses and restarts growth in 1% glucose / 1 % galactose medium.

To determine whether our allele replacement experiments recapitulated our QTL results, we fit a linear model with additive, two-way, and three-way interactions terms. We performed a likelihood ratio test to determine whether this model fit significantly better than the two-way interaction model. The coefficient of variation *R*^2^ was used to quantify the variance explained by the different QTL models. In the model with all additive and interaction terms we used a T-test to determine whether the three-way interaction term was significant.

In all of our growth experiments, we performed Welch’s t-tests to determine whether certain allele replacement strains had significantly different growth rates from each other.

To align the growth curves across a 96-well experiment for visualization purposes, for each well we identified the last time point that was less than OD 0.2, and the first time point that was greater than or equal to OD 0.2. We linearly interpolated these time-points to get an estimate for *t* at OD = 0.2 and calculated an adjusted time for all wells.

### Allele-specific expression of a hybrid diploid strain (CBS2888xBY) throughout a galactose-induction time-course

For our allele-specific RNA-sequencing experiments we mated a prototrophic BY strain (YLK1881) to CBS2888, to obtain a diploid hybrid. We performed an RNA-seq experiment for this diploid hybrid strain over a time course growth experiment where the strain was transferred from YP +2% glucose media to YP+ 2% galactose media. In more detail, we collected yeast from mid-log (OD ∼ 0.5) in glucose media. We then spun down the culture and resuspended it in galactose media (OD ∼ 0.1), we then collected yeast at 30 minutes, 1 hour, 2 hours, 4 hours, and 5.5 hours. These samples were placed in the −80 freezer for further processing. We extracted RNA from each time point using the Quick-RNA Fungal/Bacterial Kit from Zymo research. We constructed RNA-sequencing libraries using the KAPA mRNA hyperprep kits. These libraries were then sequenced on an Illumina MiSeq.

To quantify allele-specific expression (ASE) in our time course experiment, we used the WASP software to generate allele counts for each SNP site within every gene (v0.2.2)(*40*). We quantified the significance of ASE using a binomial test. For the galactose genes (*GAL1/10/7* and *GAL2*), we could not obtain ASE estimates because the reads from the CBS2888 alleles do not align to the reference. For these genes we quantified expression using Kallisto (v.0.44.0) with a reference transcriptome that contained the coding sequences of the CBS2888 galactose genes (*41*). The fold-change was calculated as the log_2_ ratio of the estimated counts of the reference and CBS2888 galactose genes.

### Curation of sequencing data used for the population and phylogenetics analysis of the galactose alleles of *S. cerevisiae*

We obtained the genome assemblies from two large collections of sequenced yeast isolates, comprising 1,277 total isolates (*12, 13*). One of these isolates had poor sequencing coverage (YCL), and we removed it from all downstream analyses. We also obtained the genome assemblies and gene annotations for the *Saccharyomyces sensu stricto* species *S. mikatae (S. mik), S. uvarum (S. uva)*, and two outgroup species *Kazaschstania africanus* (*K. afr*) and *Kazaschstania naganishii* (*K. nag*) from the yeast gene order browser (YGOB)(*42*). We obtained the genome assembly and gene annotations *S. paradoxus* (*S. par*, CBS432) from the Yeast Population Reference panel (*43*). We used the moleculo long-read assembly of CBS2888 provided by Bloom et al. as a representative strain for the alternative alleles, and the reference genome (SacCer3) as a representative strain for the reference alleles (*36*).

### Annotation of the galactose alleles from a global collection of 1,276 sequenced yeast strains

We used BLAST (v2.6.0) to align the alternative (CBS2888) and reference *GAL1, GAL10, GAL7, GAL2*, and *PGM1* promoter alleles to each of the 1,276 genome assemblies (*44*). We removed any alignments that did not cover greater than 80% of the length of the query sequence and retained the best alignment. We classified the galactose genes in each strain as alternative if the alignment had greater than 90% identity to the alternative allele and less than 90% to the reference allele. Equivalent criteria were used to classify strains as having the reference allele. The reference PGM1 promoter contained a Ty transposon, which fragmented most of the assemblies with the reference allele. For this gene, we only required that the alignment covered greater than 40% of the query sequence. Even with this relaxed criterion for some genes in some strains we were still not able to make a classification, usually due to additional assembly fragmentation. For 70 strains we were not able to assign the *PGM1* promoter, for 62 strains we were not able to assign *GAL2*, and for one strain we were not able to assign *GAL7*. In these cases, we determined which allele was present by manually combining multiple partial alignments. During this process, we identified 8 strains collected in China that had neither a reference or alternative *GAL2* or *PGM1* promoter allele. On closer inspection, all of these strains had distinct alleles (<90% sequence identity to both the reference and alternative alleles) at both of these loci.

### Analysis of the linkage disequilibrium between galactose alleles

To analyze the patterns of linkage disequilibrium (LD) between the galactose alleles, we could not use a standard measure of LD, *R*^2^, because it cannot be calculated between more than two loci. Instead, we used an entropy-based method (eLD) which generates a LD index (*ϵ*), which can be applied to arbitrary numbers of alleles (*20*). We calculated *ϵ* using the genotypes of the galactose alleles that we inferred for the 1,276 strains. Strains with the Chinese alleles were conservatively assigned as the reference. The yeast population is highly structured, and as such the null expectation for *ϵ* will be inflated relative to an unstructured population. We calculated a null distribution by randomly selecting 10,000 triplets of SNPs from different chromosomes and calculating *ϵ*.

For this analysis, we used SNPs with frequencies of between 4-5%, which is close to the observed frequency of the alternative alleles (4.4%). These SNPs were obtained by merging a VCF file provided by Peter et al. (*12*), and a VCF file we generated from data provided by Duan et al. (*13*). For the data from Peter et al. we removed sites with more than 5% missing data, indels, and sites that were not biallelic. The study by Duan et al. contained an additional 266 strains but did not provide a VCF file. We therefore downloaded the reads from the short-read archive (SRA) and generated a VCF file using the standard Genome Analysis Toolkit (GATK, v4.1.3.0) variant calling pipeline (*45*). We removed sites that had overall coverage less than 30, QUAL scores less than 100, and more than 5% missing data. We also removed sites that were indels and were not biallelic using vcftools (v0.1.15). We merged these VCF files together and set any missing sites to the reference using the “—missing-to-ref” option in bcftools (v1.9)(*46*). This merged VCF file contained a total of 1,803,186 SNPs, and had 16,451 SNPs with a frequency of between 4-5%

### Phylogenetics genetics analysis of the alternative and reference galactose alleles

To visualize the phylogenetic relationships between all 1,276 strains, we created a neighbor joining tree following similar methods as described by Peter et al. In more detail, we extracted all variants from a merged VCF file generated using bcftools with default parameters. We filtered this merged VCF to remove sites with greater than 5% missing data and sites with a minor allele frequency less than 5%. Next we generated a dissimilarity matrix using SNPRelate (v1.18.1)(*47*). This matrix was used to build a neighbor joining tree with ape (v5.3)(*48*). We visualized this tree using the ggtree (v1.16.4) package (*49*). We colored the branches based on which combination of the galactose alleles each strain had.

To visualize the phylogeny of the alternative galactose alleles, we performed phylogenetic clustering of the *GAL1, GAL10, GAL7* coding sequences from CBS2888 (alternative), SacCer3 (reference), S. par, S. mik, S. uva, K. afr, and K. kna(*35, 50*). We did not use *GAL2* in this analysis because this gene is only found in the *Saccharyomyces sensu stricto.* We aligned each gene using Clustal Omega (v1.2.4) and concatenated the sequences. We used RAxML (v 8.1.21) with a GTR + GAMMA model for tree generation (*51*). We set the outgroups to be K. afr and K. kna. We performed 100 rapid bootstraps to assess the confidence of the branches.

To investigate the phylogenetic relationships between the alternative, Chinese, and reference *GAL2* genes, we performed phylogenetic clustering of the *GAL2* coding sequence from CBS2888 (alternative), BAM (Chinese), SacCer3 (reference), S. par, S. mik, S. uva. *GAL2* is only found in the *Saccharyomyces sensu stricto*, so we used the K. afr and K. nag *HXT7* genes as the outgroups. We aligned the genes and performed phylogenetic clustering using RAxML using the options described above. We annotated the cytosolic, extracellular, and trans-membrane domains of the S. cer *GAL2* gene using TMHMM (*52*). We aligned the DNA sequences, extracted the aligned sequences of each domain, and performed phylogenetic clustering using RAxML. We added the species *S. eubayanus* to this analysis because it has a duplicated *GAL2.* We also performed sub-clustering phylogenetic analysis using the protein sequences of the N-terminal domain comprising the first 67 amino acids.

### Population genetics analysis of the alternative and reference galactose alleles

We estimated the synonymous substitutions per site (dS) between the reference and alternative galactose genes using the codonseq package of Biopython (v1.70) with the NG86 method (*53, 54*). We note that this method utilizes the Jukes-Cantor correction that explicitly incorporates back mutations and converts the raw proportion of observed synonymous changes into a time linear distance between the sequences. If there are many differences between the sequences this can lead to estimates of dS that are greater than 1. To calculate a 95% confidence interval on all our dS estimates, we performed 200 bootstraps in which we resampled codons with replacement and then recalculated the dS. When the number of synonymous substitutions is too large the NG86 method can return an error, for these bootstrap samples dS was set to 3.23. To obtain a background distribution of synonymous differences, we performed a global alignment using ssearch36 (v36.3.8f) to identify the reference genome open-reading frames in the CBS2888 genome (*55*). We only considered alignments that contained a complete open reading frame. We calculated the dS for each aligned gene using codonseq. We aligned the alternative and reference *PGM1* promoters and calculated the distance between the sequences using the DistanceCalculator from Biopython with the ‘identity’ method. For every pair of neighboring genes in the reference genome, if such pairs both align to the same CBS2888 contig we extracted the sequences between them for both the reference and CBS2888 assemblies and calculated percent identity using the method described above for the *PGM1* promoters.

To estimate the number of generations since *S. cerevisiae* split from *S. paradoxus, S. mikatae*, and *S. uvarum* (*56*), we calculated the dS for a random sample of 161 genes that were similar in length to the galactose genes. These alignments were concatenated, and we calculated the dS using codonseq.

### Analysis of synonymous substitutions per site in CBS2888

Using the aligned genes between CBS2888 and the reference genome, we calculated the dS in 200 amino acid overlapping windows genome-wide using codonseq. We used a step of 10 amino-acids for the windows.

## Acknowledgements

We thank Olga T Schubert, Eyal Ben-David, Longhua Guo, and Stefan Zdraljevic for helpful manuscript feedback and edits.

## Author contributions

JB performed experiments with assistance from MS and JSB. JB analyzed data with assistance from JSB. LK and JSB supervised the project. JB, JSB, and LK wrote the manuscript. All authors discussed and agreed on the final version of the manuscript.

## Data and materials availability

Sequencing data are available under NCBI BioProject PRJNA575066.

