## Supplementary Notes and Figures for "Ancient balancing selection maintains incompatible versions of a conserved metabolic pathway in yeast"

**Supplementary Note 1**

**Estimating the age of the balancing selection by dating the linked neutral sites.**

Estimates of divergence at linked neutral sites can be used to provide a lower bound on the age of the balancing selection. To date the divergence between the alternative and reference neutral linked sites, we combined the estimated number of synonymous substitutions per site with the measured mutation rate of $\mu = 3.8 \cdot 10^{-10}$ per site per generation in yeast. Under neutral theory, the dS value of 1.20 (dS=1.20, 95% C.I.=0.81-2.10) for the neutral site near *KAP104* corresponds to a split between this reference and alternative site approximately 1.58 billion generations ago. *S. cerevisiae* split from *S. paradoxus* ~0.61, *S. mikatae* ~1.17, and *S. uvarum* ~1.91 billion generations ago which suggests that this balancing selection has been acting on these galactose alleles since at least the time of the *S. mikatae* and *S. cerevisiae* split.

**Supplementary Note 2**

**The alternative and Chinese galactose alleles provide additional evidence that the balancing selection is ancient.**

In our global analysis of the galactose alleles, we identified distinct *GAL2* alleles in strains isolated from China. These alleles contain *GAL2* genes duplicated in tandem, and *GAL2* is also duplicated in two other yeast species, *S. uvarum* and *S. eubayanus*. The centromere-proximal *GAL2* paralog is denoted as *GAL2a* and the other *GAL2b*. We phylogenetically clustered the *GAL2* alleles and found that the Chinese and diverged *GAL2* alleles are more closely related to each other than to any other *GAL2* alleles (Supplementary Figure 14). In contrast, when we clustered the DNA sequences of the cytosolic N-terminal domains, we found that the N-termini of the diverged and reference *GAL2b* genes are more similar to each other than the N-termini of their *GAL2a* genes (Supplementary Figure 15). We further clustered the protein sequences from the N-terminal domains, which revealed that all the *GAL2b* alleles are more closely related to each other than to any other *GAL2* alleles. This suggests that both paralogs have conserved and distinct roles in *S. cerevisiae*, *S. uvarum*, and *S. eubayanus*.

The most parsimonious explanation for the origin of the alternative alleles is that they arose in an ancestor of *S. uvarum*, *S. eubayanus*, and *S. cerevisiae*, and have been maintained by balancing selection ever since. However, our population genetics analysis cannot rule out a later introgression into an ancestor of *S. cerevisiae*. The Chinese and alternative *GAL2* alleles diverged from each other approximately 1.6 billion generations ago. These alleles also phylogenetically cluster and form an outgroup to the *Saccharyomyces sensu stricto* species complex (Supplementary Figure 14). These observations provide us with another explanation for the origin of the alternative and Chinese galactose alleles, that if these alleles were originally introgressed, it happened over 1.6 billion generations ago.

### Supplementary Note 3

#### Phylogenetics and population genetics analyses of the alternative galactose alleles in a global sample of 1,276 sequenced yeast strains.

We used the genomes of representative strains for the phylogenetics and population genetics analyses that we presented in the main text. In particular, we used the CBS2888 genome for the alternative alleles (*16*), the BAM genome for the Chinese alleles (*35*), and the reference genome (SacCer3) for the reference alleles (*36*). To ensure that these genomes were representative of strains with the alternative, Chinese, and reference alleles, we repeated all our population genetics and phylogenetics analyses in the genome assemblies from the global collection of 1,276 sequenced yeast strains (*12*, *13*).

Using a custom script, we extracted as many genes as possible from the 1,276 genome assemblies (Methods). We calculated the number of synonymous substitutions per site in 200 amino acid windows with a 10 amino acid step genome-wide when compared to the reference genome. In agreement with our analysis of the CBS2888 genome, we observed a strong signature of balancing selection at the galactose genes of strains classified as having the alternative or Chinese galactose alleles (Supplementary Figure 16). We did not observe this signature when comparing strains classified as having the reference galactose alleles.

We extracted *GAL1*, *GAL10*, and *GAL7* alleles from the genomes of 1,234 of the 1,276 strains with complete assembled ORFs at these sites. We removed the 3 strains with reference alleles at all loci except *GAL7*. We performed phylogenetic clustering of the genes from the remaining 1,231 strains and verified that all of the alternative *GAL1/10/7* alleles form an outgroup to the *Saccharyomyces sensu stricto* species complex (Supplementary Figure 17).

We calculated the synonymous substitutions per site between the reference genome and all *GAL1*, *GAL10*, and *GAL7* genes (Supplementary Figure 18). In agreement with our comparison between CBS2888 and reference galactose alleles, the number of synonymous substitutions per site was high in strains classified as having the alternative alleles (*GAL7* dS=2.50, range=0.94-2.88; *GAL10* dS=2.64, range=2.53-2.85; *GAL1* dS=2.13, range=2.05-2.52). The number of synonymous substitutions per site was low when comparing the alleles of strains that we annotated as having the reference alleles (*GAL7* dS=0.01, range=0-0.26, *GAL10* dS=0.01, range=0-0.016, *GAL1*=0.01, range=0-0.041)

**Supplementary Note 4**

**The alternative and Chinese galactose alleles fall outside the *Saccharyomyces sensu stricto* species complex.**

The Chinese and alternative *GAL2* alleles both contain a *GAL2* duplication. We compared the centromere-proximal alternative and Chinese *GAL2* alleles and found that these two alleles are more similar to each other (dS=1.37) than they are to the reference (Chinese *GAL2* compared to reference dS=1.91, alternative GAL2 compared to reference dS=2.38). Phylogenetic clustering revealed that the Chinese and alternative *GAL2* genes cluster with each and fall outside the *Saccharomyces sensu stricto* (Supplementary Figure 14).

The Chinese and alternative *PGM1* promoter alleles are more similar to each other (70.6% sequence identity) than either is to the reference (Chinese *PGM1* promoter compared to reference, sequence identity=47.0%; alternative *PGM1* promoter compared to reference, sequence identity=46.7%). The low sequence conservation between the *PGM1* promoters in outgroup species prevented us from performing phylogenetic clustering, however, we did note that the alternative and Chinese *PGM1* promoter alleles are missing a lysine tRNA that is present in every species of the *Saccharomyces sensu stricto* (Supplementary

Figure 19). This places the Chinese and alternative *PGM1* promoter alleles outside the *Saccharomyces sensu stricto*.

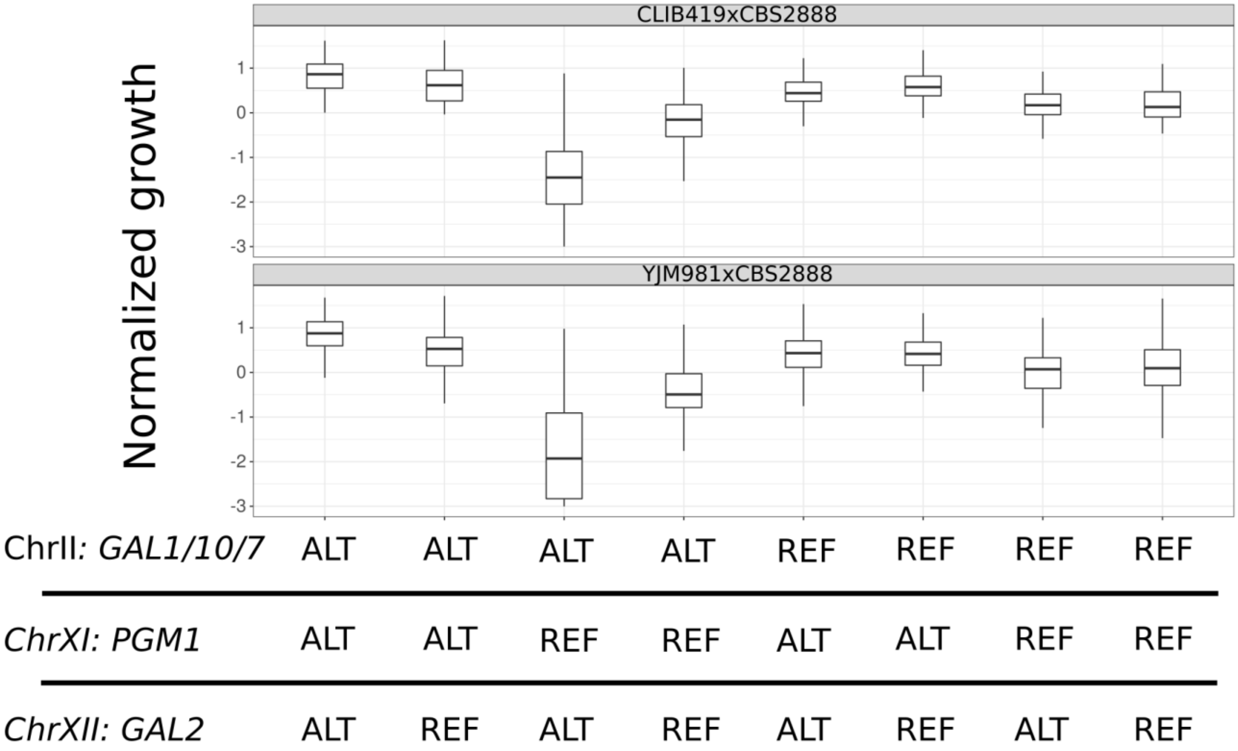


**Supplementary Figure 1. Replication of the three-way genetic interaction another panel of segregants.**

top) Boxplots showing the growth (s.d units) of 867 segregants on 2% galactose agar plates. These segregants were derived from a cross between CBS2888 and YJM981. bottom) Boxplots showing the growth of 943 segregants on 2% galactose agar plates. These segregants were derived from a cross between CBS2888 and CLIB219. Segregants are partitioned on the x-axis based on their genotypes at eight combinations of the three galactose loci (ChrII: *GAL1/10/7*, ChrXI*: PGM1*, and ChrXII: *GAL2*). Alleles at the QTL loci from CBS2888 are designated as ALT, and alleles from CLIB219 and YJM981 are designated as REF.

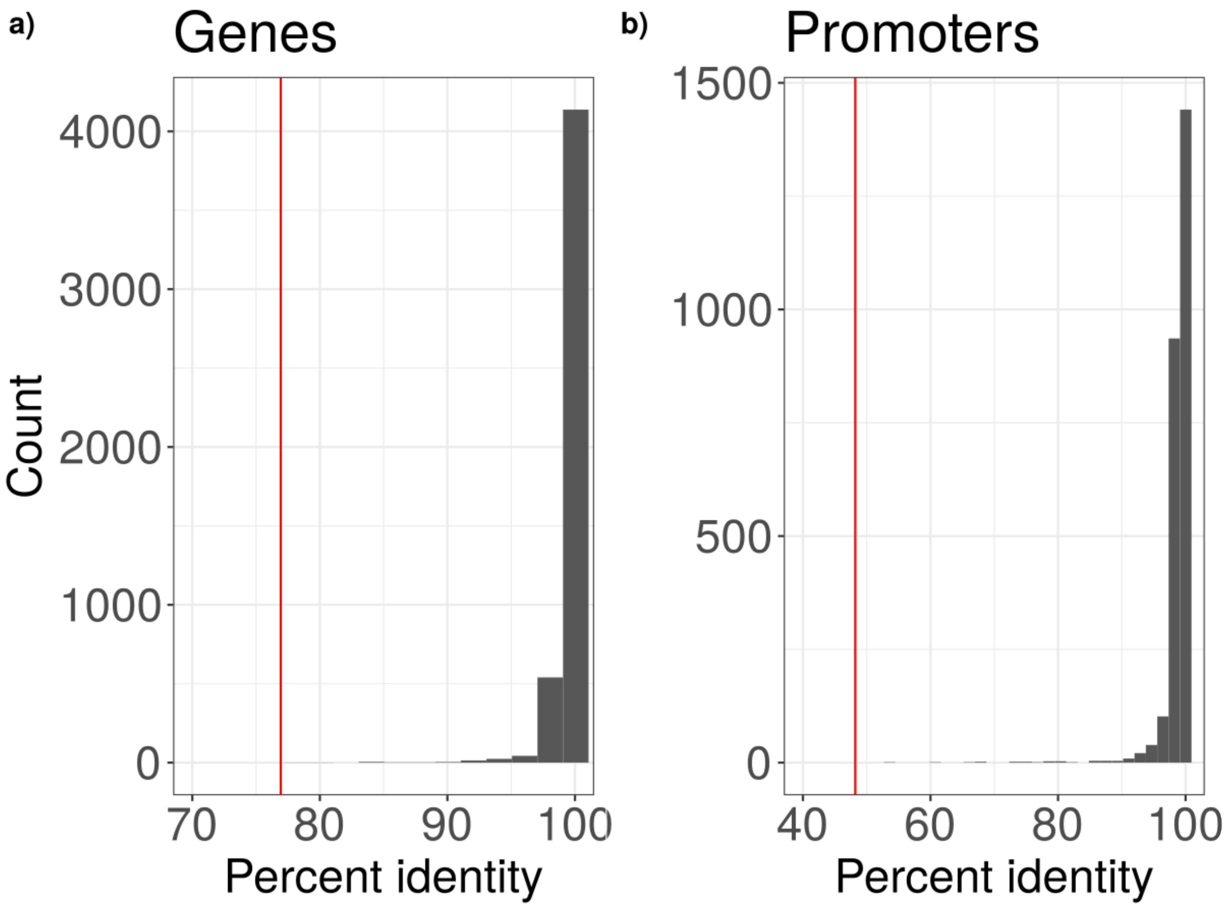


**Supplementary Figure 2: Sequence identity between the genes and promoters of the reference and CBS2888.**
a) Histogram of the percent identity of 4,780 genes in the CBS2888 genome when these genes were aligned to the reference genes. The average sequence identity of the diverged *GAL1*, *GAL10*, *GAL7*, and *GAL2* (77%) when aligned to the reference galactose alleles is shown in red. b) Histogram of the percent identity of 2,583 promoters of CBS2888 when aligned to the reference. The sequence identity of the diverged PGM1 promoter (48%) when aligned to the reference promoter is shown in red.

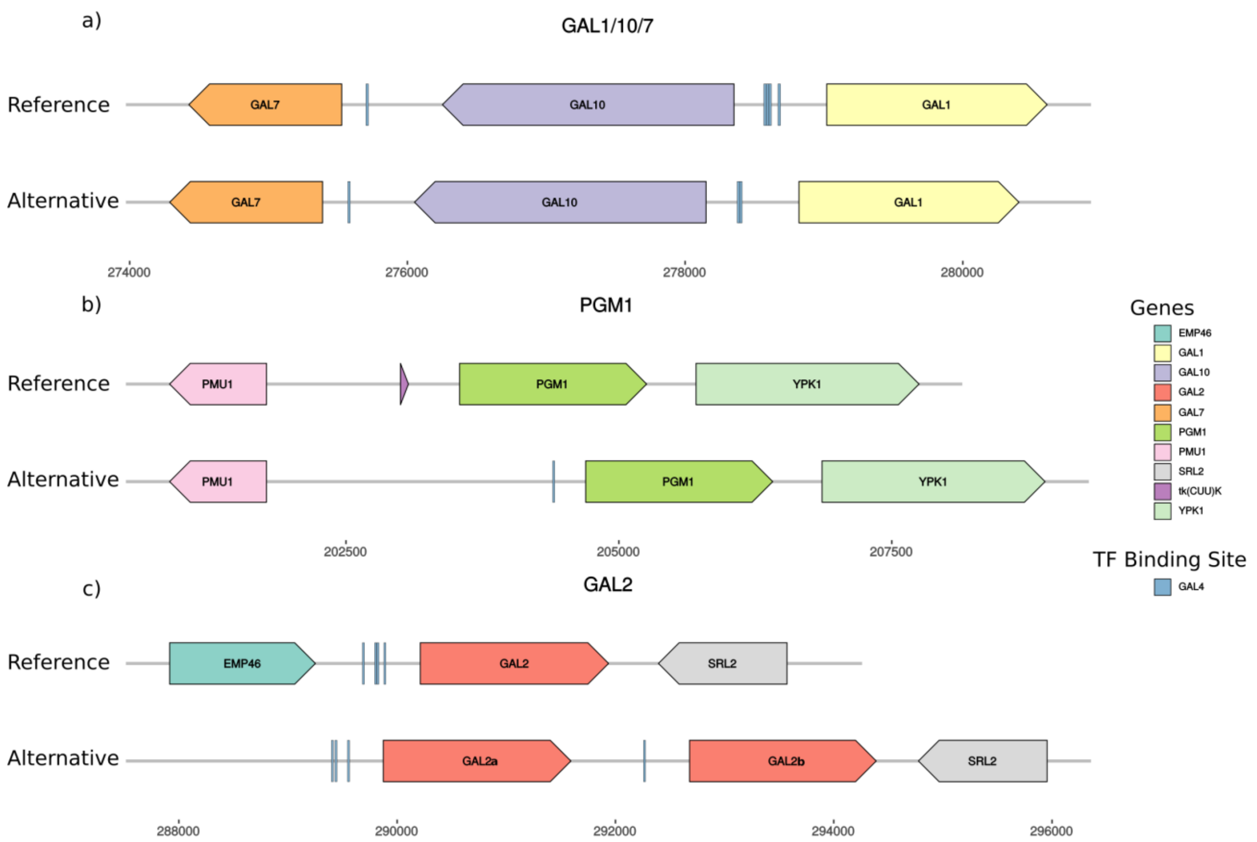


**Supplementary Figure 3: Genomic organization of the alternative and reference galactose alleles.**

Regional genome plots showing the relative lengths and locations of the genes flanking the galactose alleles. We also annotated any *GAL4* upstream activating sequence (UAS) that we found at each of the loci. a) *GAL1/10/7*, b) *PGM1*, c) *GAL2*. The centromere-proximal copy of *GAL2* in CBS2888 (alternative) is denoted as *GAL2a*, and the other copy as *GAL2b*.

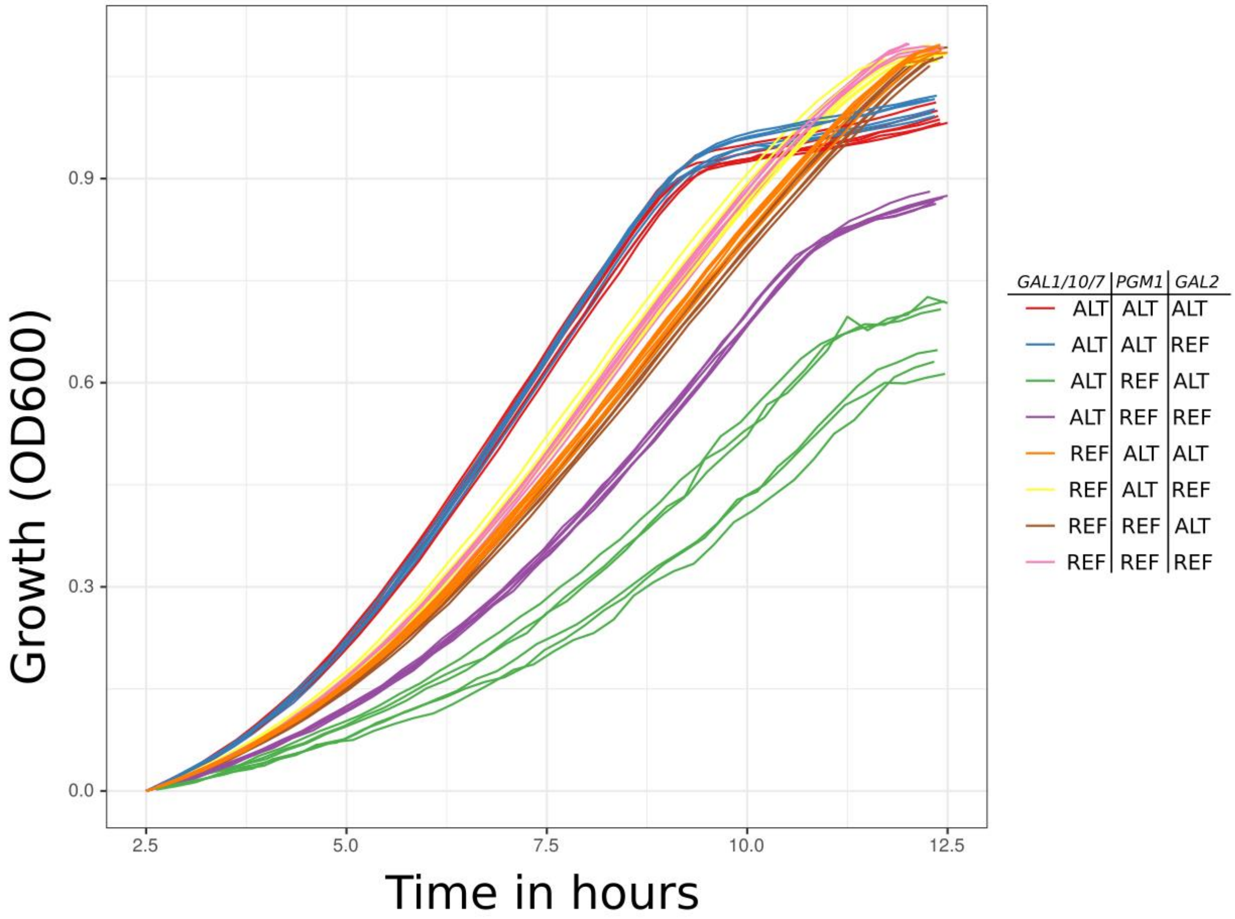


**Supplementary Figure 4: Growth curves for allele replacement strains that contain all eight combinations of the alternative and reference galactose alleles.**

Growth curves for each allele replacement strain. Each growth curve is colored according to the genotype of the corresponding strain.

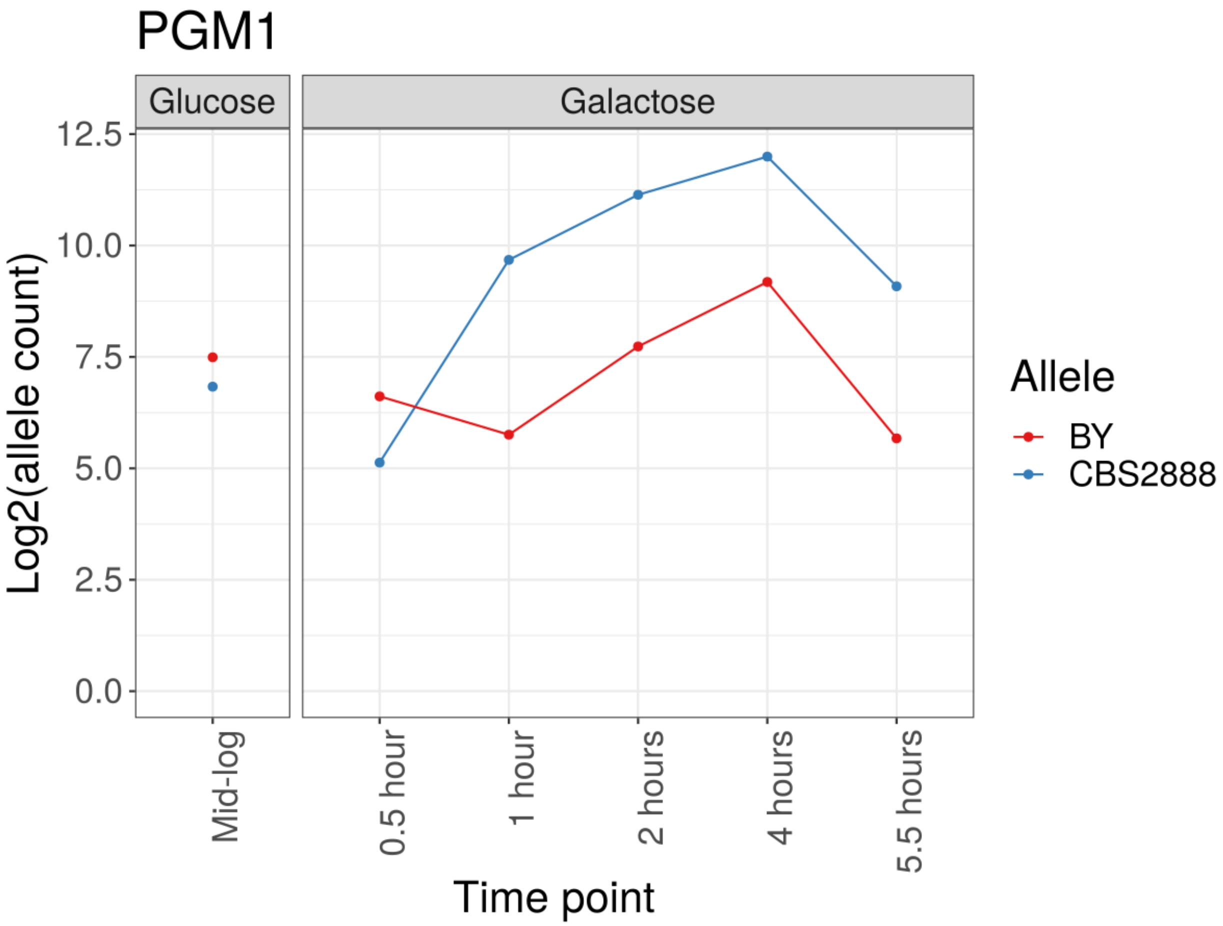


**Supplementary Figure 5: Allele-specific expression of PGM1 throughout a galactose induction time course.**

Allele-specific expression of a hybrid (CBS2888xBY) that is heterozygous for the alternative and reference alleles when grown in 2% glucose medium and transferred to 2% galactose medium. The line graph shows the $\log_2$ allele counts for the CBS2888 (alternative) alleles and BY (reference) alleles.

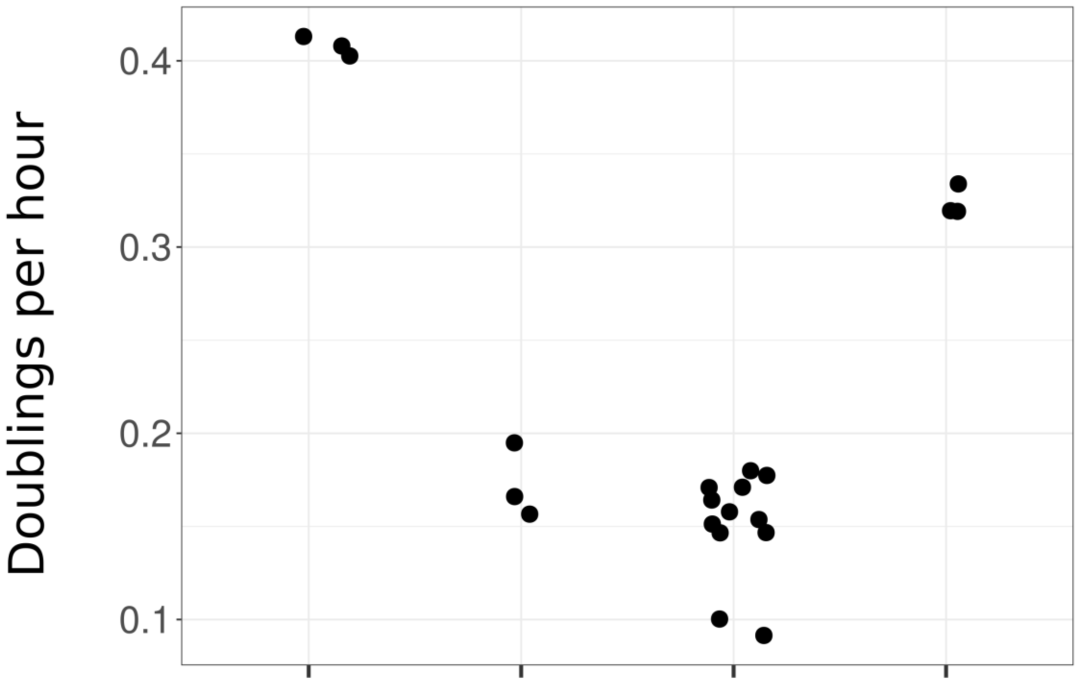


| | | | | |
|---|---|---|---|---|
| *ChrII:GAL1/10/7* | ALT | ALT | ALT | REF |
| *ChrXI: PGM1* | ALT | REF | ALT | REF |
| *GAL4-UAS* | WT | N.A. | Δ GAL4-UAS | N.A. |
| *ChrXII: GAL2* | ALT | ALT | ALT | REF |

**Supplementary Figure 6: Galactose responsive *PGM1* is necessary for the proper functioning of the alternative galactose pathway.**

Growth rates of allele replacement strains when grown in 2% galactose medium. The strains are from left to right: A strain with all three alternative alleles. A strain with the reference promoter and the alternative *GAL1/10/7* and *GAL2*. A strain with all three alternative alleles and a non-functional GAL4-UAS ($CGGN_{11}CCG$-> $CGGN_{11}ACG$). A strain with all three reference alleles. Each dot shows a biological replicate.

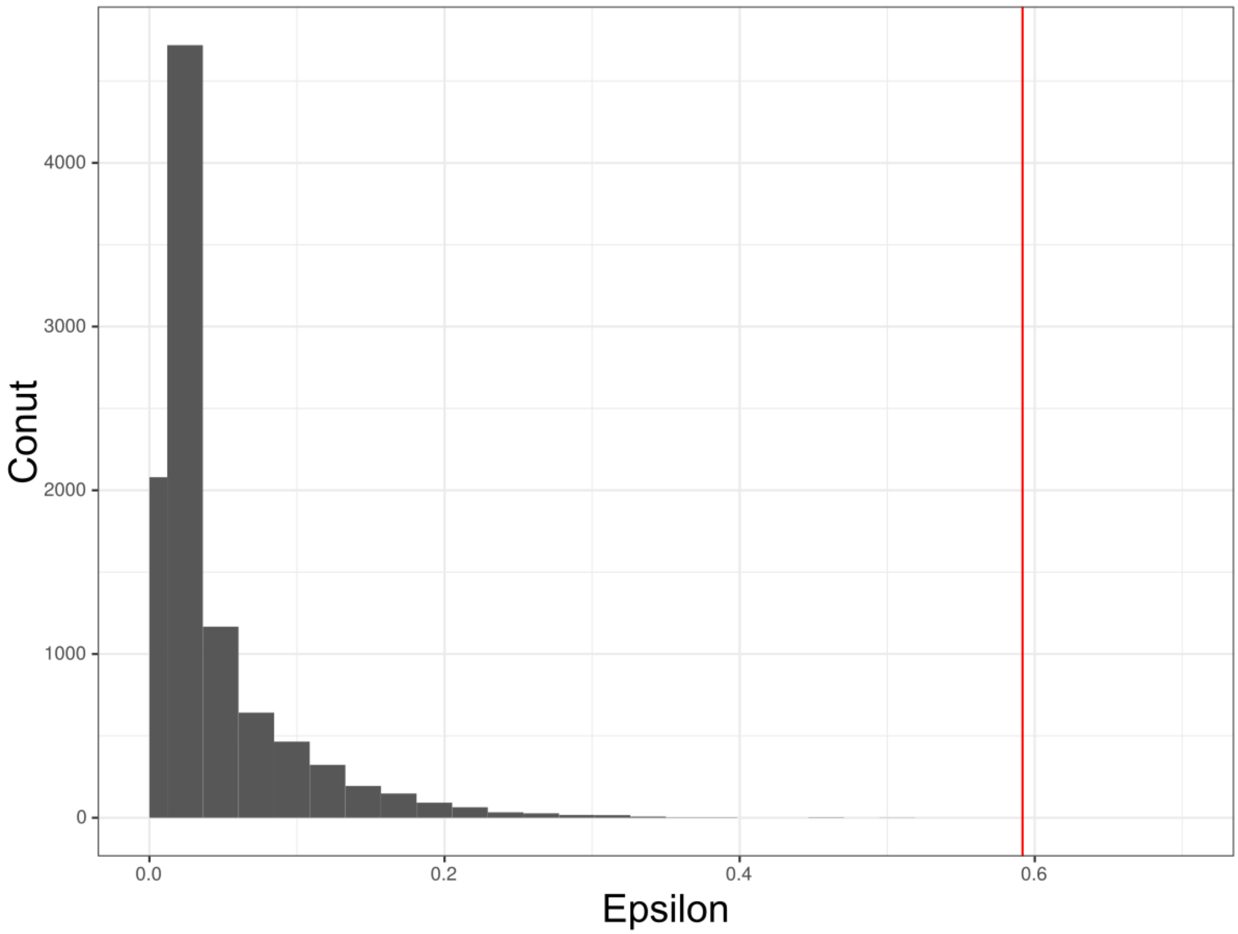


**Supplementary Figure 7: Linkage disequilibrium between 10,000 random sets of three SNPs.**

Distribution of the entropy-based linkage disequilibrium index ($\epsilon$) of 10,000 random sets of three SNPs with a similar frequency (4-5%) to the diverged galactose alleles. In red is the value of $\epsilon$ of the galactose alleles (0.592). The largest value we observed in our 10,000 permutations was 0.50. A larger $\epsilon$ indicates that the sites are in high linkage disequilibrium.

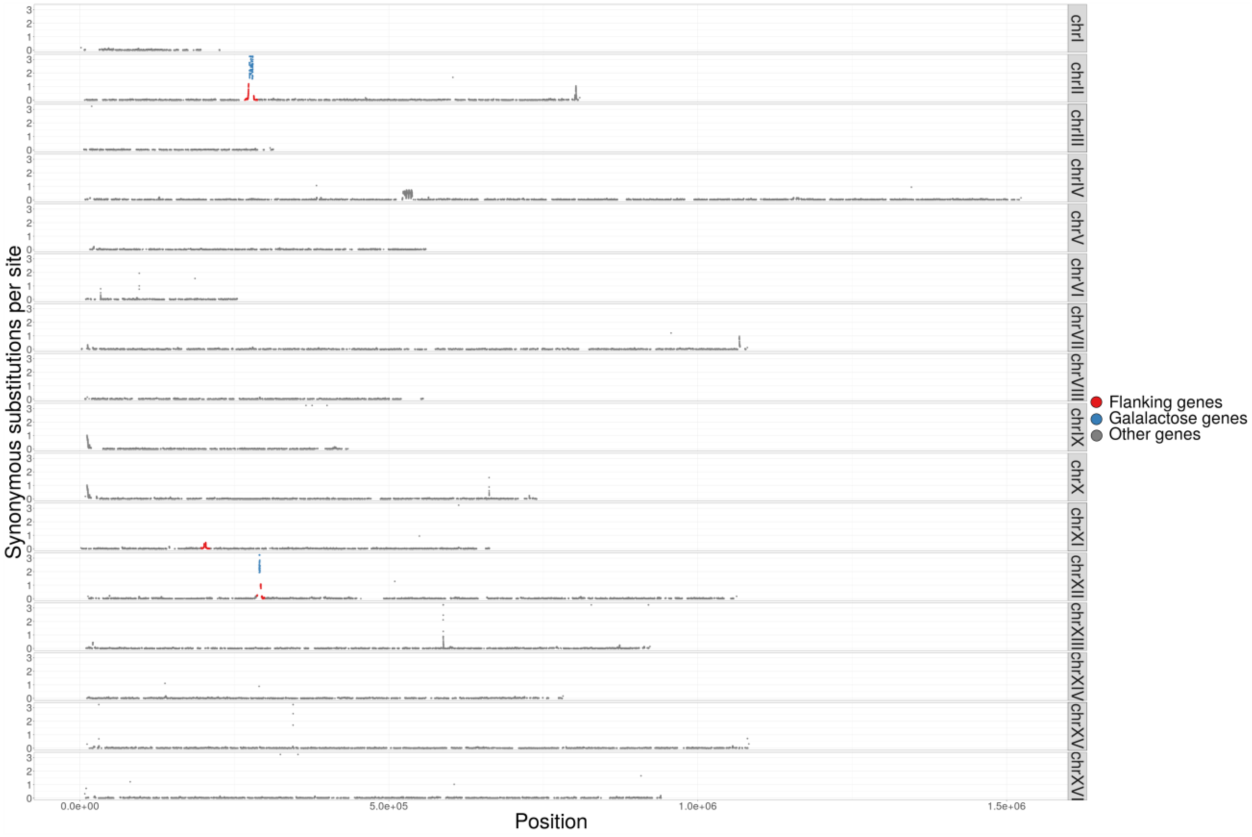


**Supplementary Figure 8: Genome-wide distribution of the synonymous substitutions per site (dS) 200-amino acid sliding windows between the CBS2888 and SacCer3 genes.**

The distribution of dS in 200-amino acid sliding windows with a step of 10-amino acids when the CBS2888 genes are aligned to the SacCer3 genes. We have highlighted in red genes near the galactose loci, in blue are the galactose genes (*GAL1*, *GAL10*, *GAL7*, and *GAL2*), and in grey are all other genes. *GAL1/10/7* are on ChrII, *PGM1* is on ChrXI, and *GAL2* is on ChrXII.

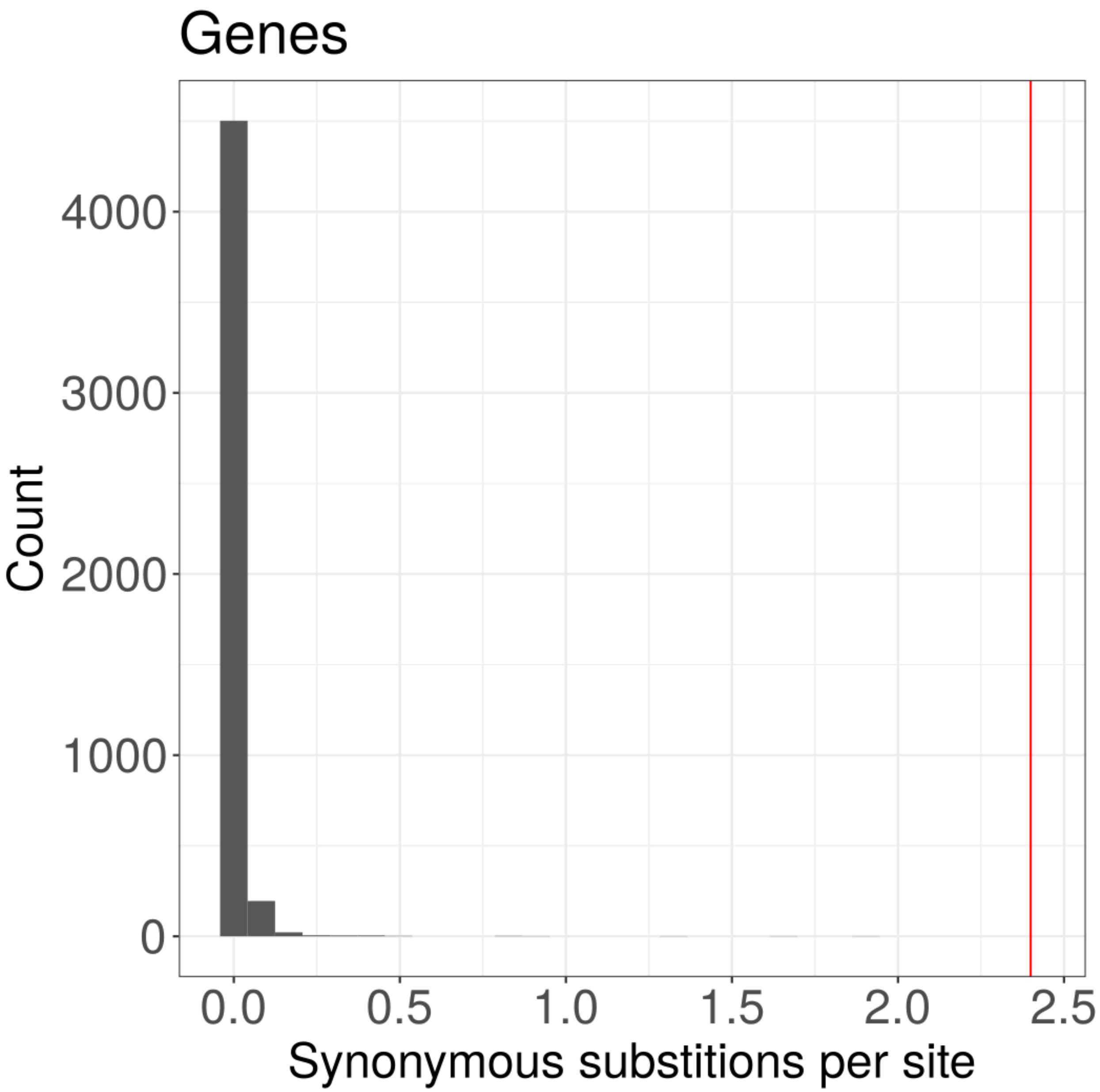


**Supplementary Figure 9: Synonymous substitutions per site (dS) between the genes of SacCer3 and CBS2888.**

Histogram of the dS of the CBS2888 genes when compared to the reference genes. In red is shown the average dS of the CBS2888 diverged *GAL1*, *GAL7*, *GAL10*, and *GAL2* galactose alleles when compared to the reference genes.

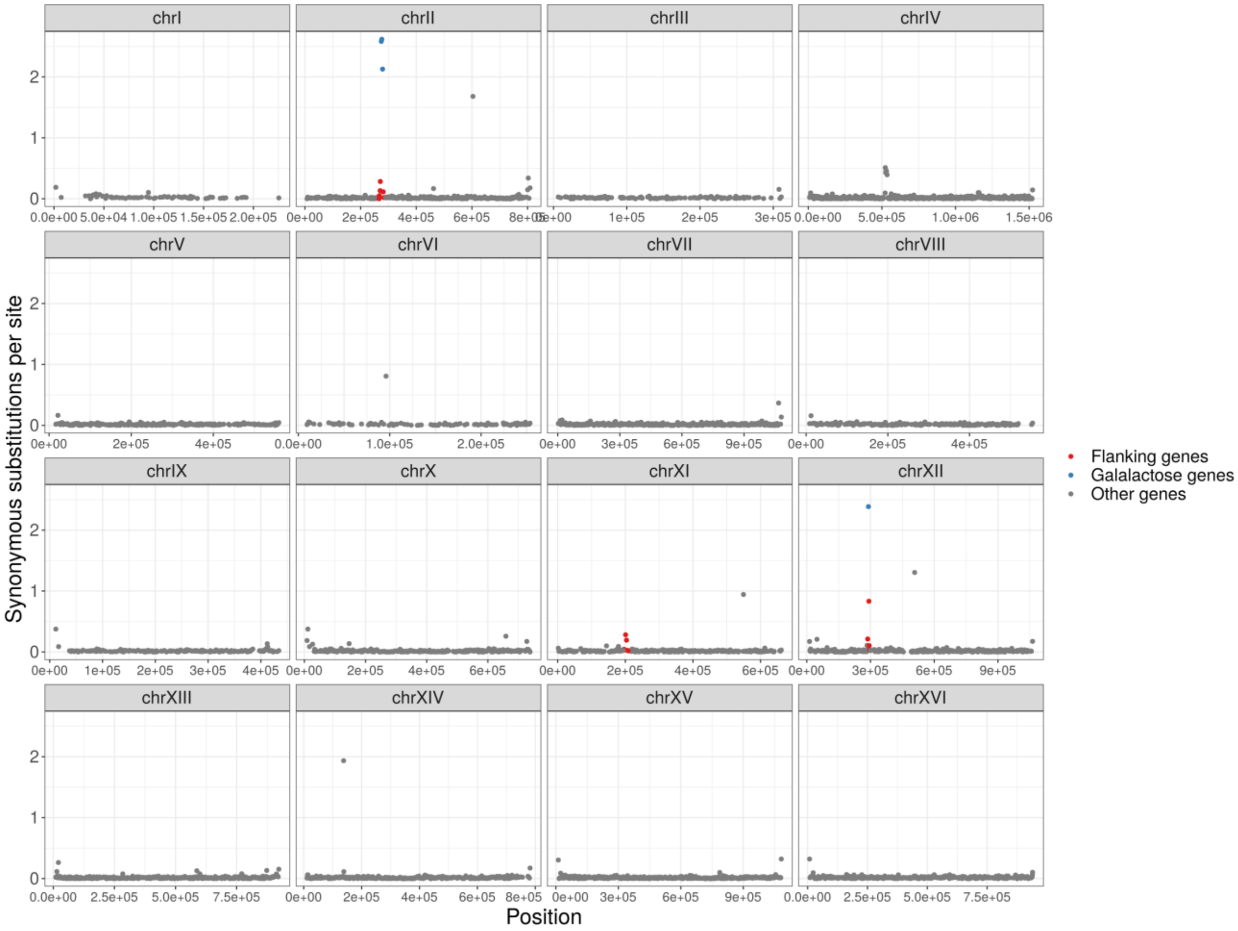


**Supplementary Figure 10: Genome-wide distribution of the synonymous substitutions per site (dS) between the genes of SacCer3 and CBS2888.**

The distribution of the dS of the CBS2888 genes when they are aligned to the reference genes. We have highlighted in red genes near the galactose loci, in blue are the galactose genes (*GAL1*, *GAL10*, *GAL7*, *GAL2*), and in grey are all other genes. *GAL1/10/7* are on ChrII, *PGM1* is on ChrXI, and *GAL2* is on ChrXII.

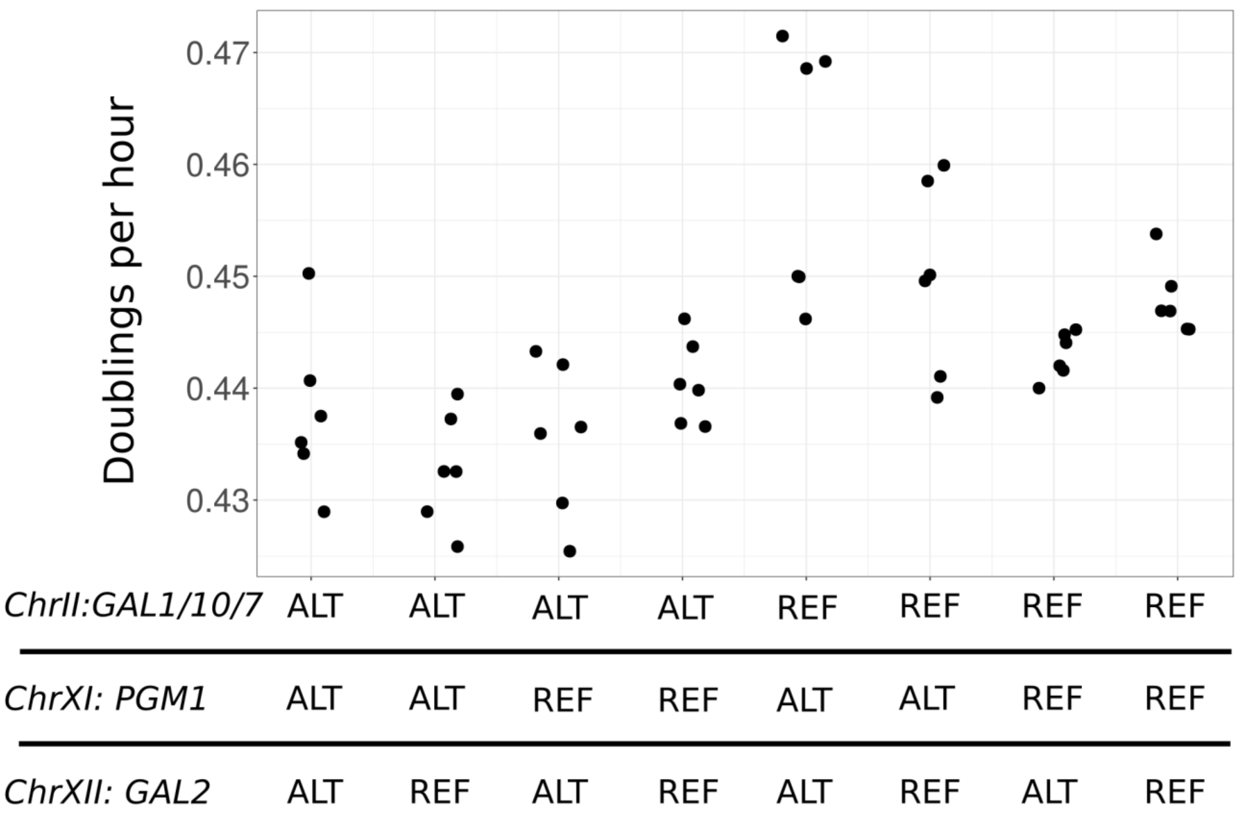


**Supplementary Figure 11: Growth of allele replacement strains in 2% glucose medium.**

The doublings per hour in 2% glucose medium for allele replacement strains. Alleles at the QTL loci from CBS2888 are designated as ALT, and alleles from BY are designated as REF. Each dot shows a biological replicate.

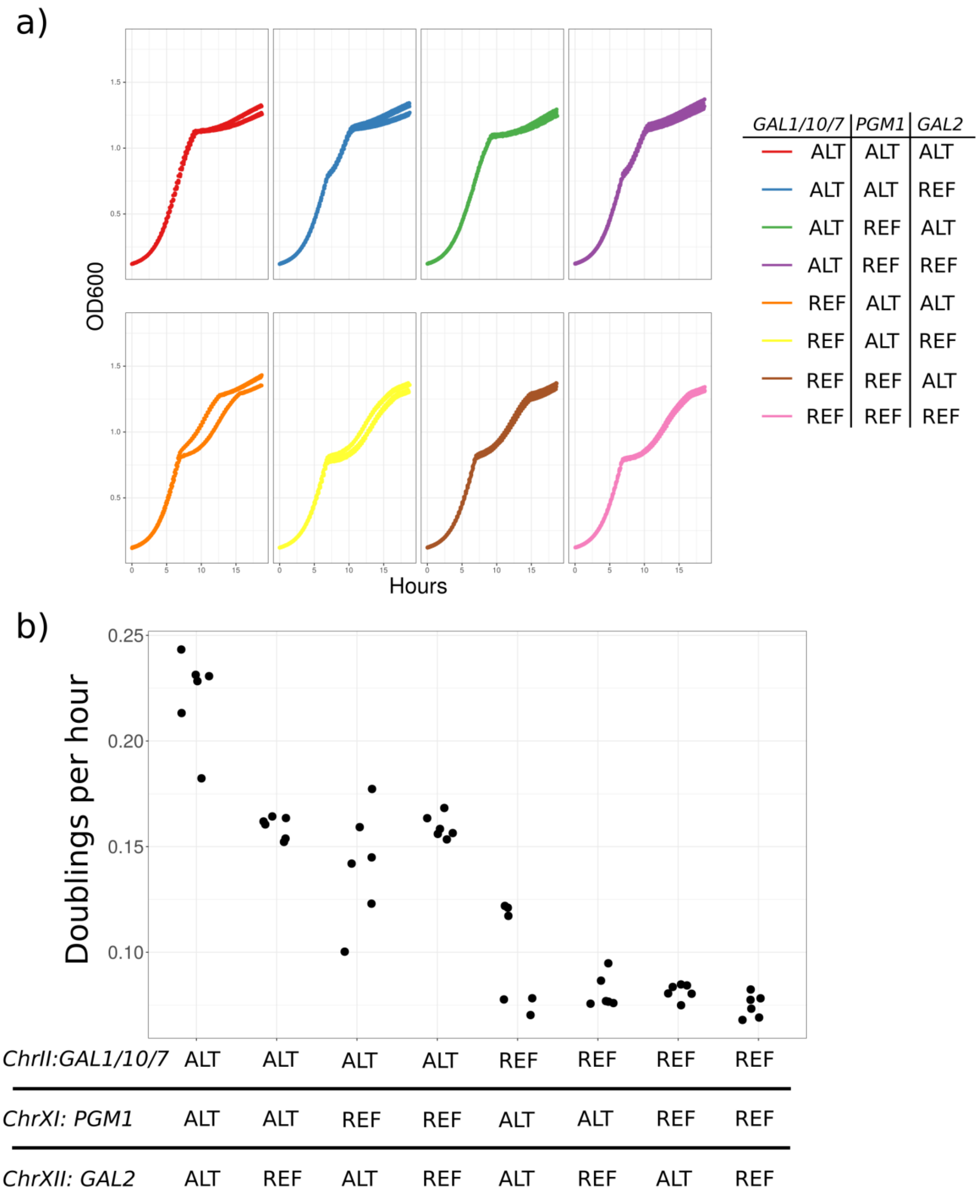

a)
OD600
Hours
GAL1/10/7 | PGM1 | GAL2
ALT ALT ALT
ALT ALT REF
ALT REF ALT
ALT REF REF
REF ALT ALT
REF ALT REF
REF REF ALT
REF REF REF
b)
Doublings per hour
0.25
0.20
0.15
0.10
ChrII:GAL1/10/7 ALT ALT ALT ALT REF REF REF REF
ChrXI: PGM1 ALT ALT REF REF ALT ALT REF REF
ChrXII: GAL2 ALT REF ALT REF ALT REF ALT REF

**Supplementary Figure 12: Strains with the diverged galactose alleles do not experience a diauxic shift when they switch from glucose to galactose medium.**

a) Growth curves for all eight allele replacement strains when growth in 1% glucose/1% galactose medium. Each growth curve is a summary of three biological replicates from each of the two strains for all eight possible combinations of alleles. b) The growth rate between 0.8 and 1.1 OD600 units for all eight allele replacement strains. Each dot represents a biological replicate. Quantifying this range of the growth curve captures the degree of diauxic shift.

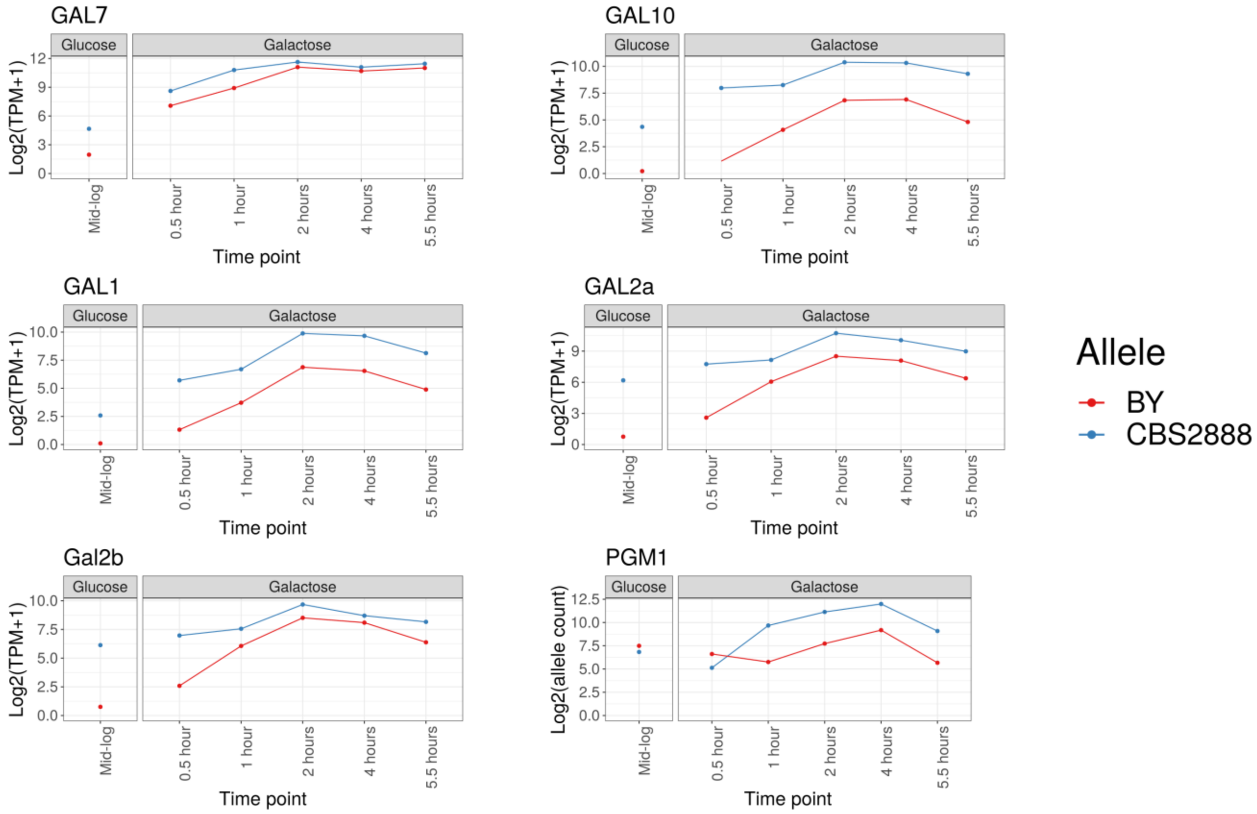


**Supplementary Figure 13: The alternative (CBS2888) galactose alleles are constitutively expressed in glucose and are more upregulated in galactose.**

Allele-specific expression of a hybrid (CBS2888xYJM981) that is heterozygous for the alternative and reference alleles when grown in 2% glucose medium and transferred to 2% galactose medium. The line graph shows the $\log_2$ allele counts or transcripts per millions (TPM) for the CBS2888 (alternative) alleles and BY (reference) alleles. We denote the centromere-proximal copy of *GAL2* in CBS2888 as *GAL2a*, and the other one as *GAL2b*.

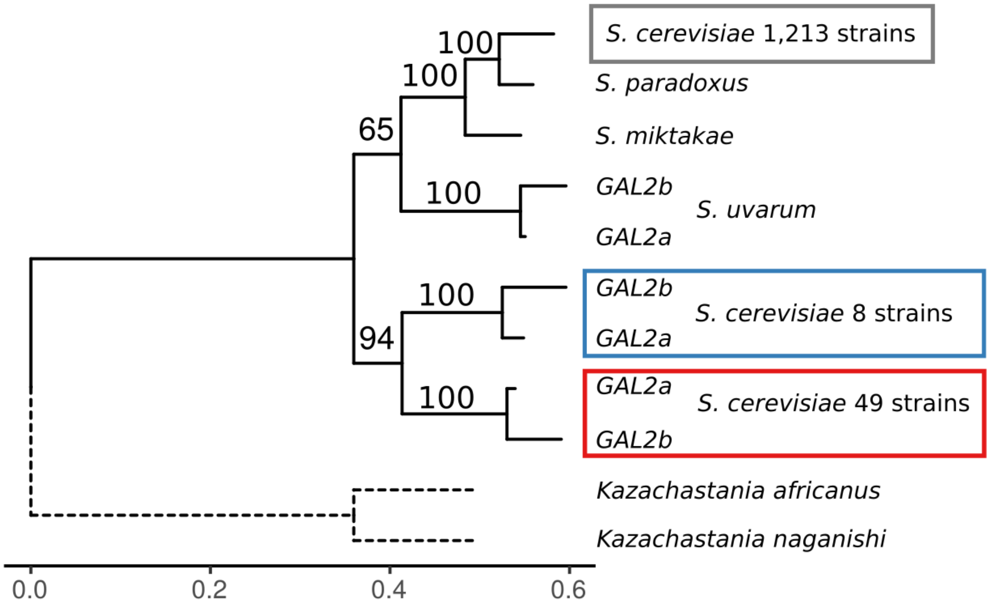


**Supplementary Figure 14: Phylogenetic clustering of the *GAL2* alleles.** Bootstrapped maximum likelihood phylogenetic trees of the *GAL2* alleles. Gene sequences were extracted from the genomes of CBS2888 (alternative), SacCer3 (reference), other members of the *Saccharoymyces sensu stricto* (*S. uvarum*, *S. mitakae*, *S. paradoxus*), and the outgroup species (*Kazachstania africanus* and *Kazachstania naganashii*). The outgroup branches (dotted lines) were rescaled to the average branch length. The centromere-proximal copy is denoted as *GAL2a*, and the other copy as *GAL2b*. The *S. cerevisiae* alleles are colored according to their classification; alternative (red), Chinese (blue), and reference (grey). Scale bar shows the estimated number of substitutions per site.

a) N-terminal DNA sequence

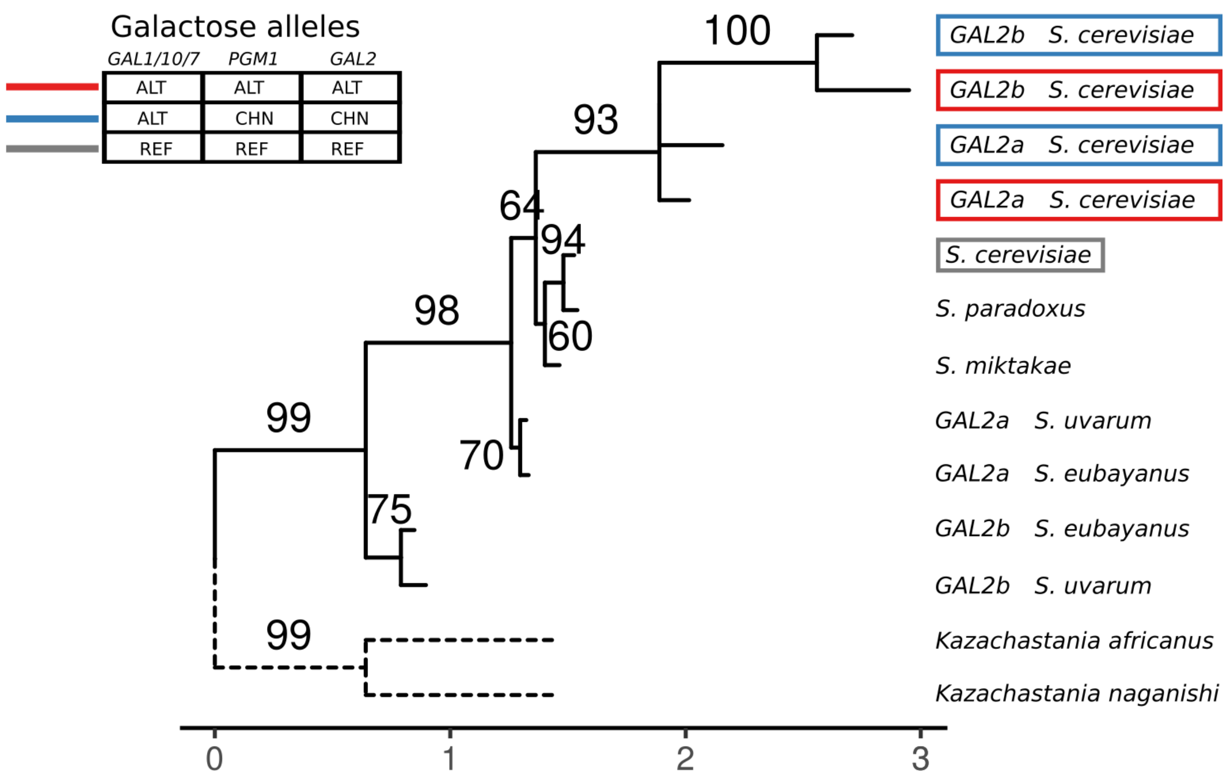


b) N-terminal Protein sequence

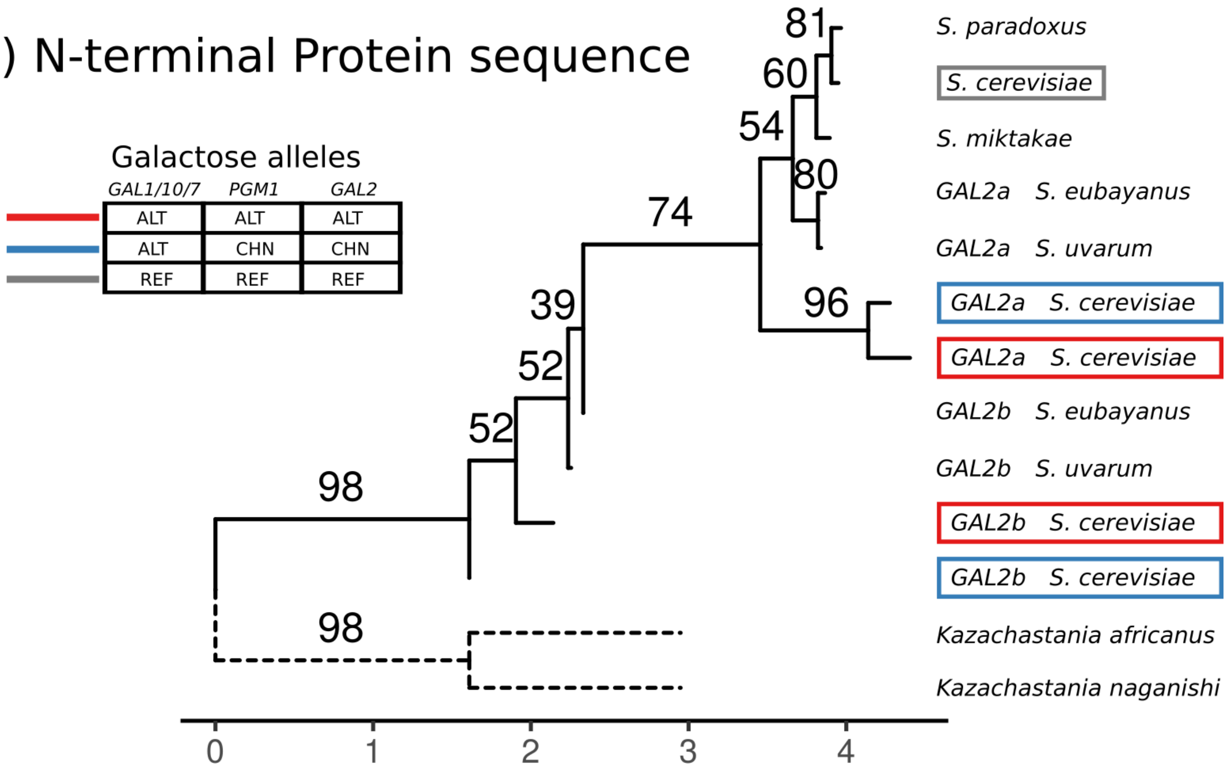

**Supplementary Figure 15: The cytosolic N-terminal regions of the GAL2a and GAL2b genes are distinct and have been conserved for millions of years.**

Bootstrapped maximum likelihood trees of the cytosolic N-terminal region of the *GAL2* genes. Gene sequences were extracted from the genomes of CBS2888 (alternative), SacCer3 (reference), other members of the *Saccharoymyces sensu stricto* (*S. uvarum, S.eubayanus, S. mitakae, S. paradoxus*), and the outgroup species (*Kazachstania africanus* and *Kazachstania naganashii*). The diverged and Chinese *GAL2* genes duplicated in tandem, and this duplication is also found in *S. uvarum* and its sister species *S. eubayanus*. The *S. cerevisiae* alleles are colored according to their classification; alternative (red), Chinese (blue), and reference (grey). The centromere-proximal *GAL2* paralog is a denoted as *GAL2a* and the other copy as *GAL2b*. Scale bar shows the estimated number of substitutions per site.

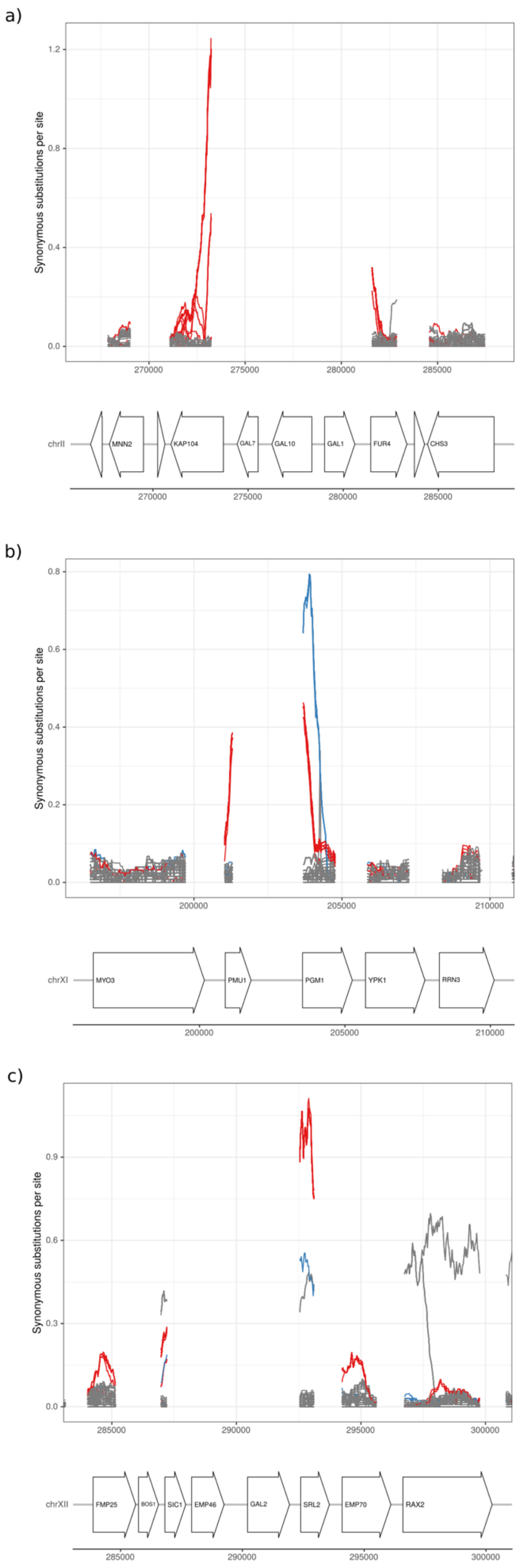


**Supplementary Figure 16: A signature of balancing selection in the global yeast population.**

Synonymous substitutions per site (dS) in 200-amino acid sliding windows between the 1,276 sequenced yeast strains and the reference in regions surrounding the galactose loci. The dS for strains with only alternative alleles are shown in red. The dS for strains with only reference alleles are shown in grey. The dS for strains from China with the alternative *GAL1/10/7* allele and alleles of *GAL2* and the *PGM1* promoter that differ from both the reference and the alternative alleles are shown in blue. The dS was not estimated for *EMP46* due to the presence of an early stop codon interrupting this gene in strains with the alternative alleles.

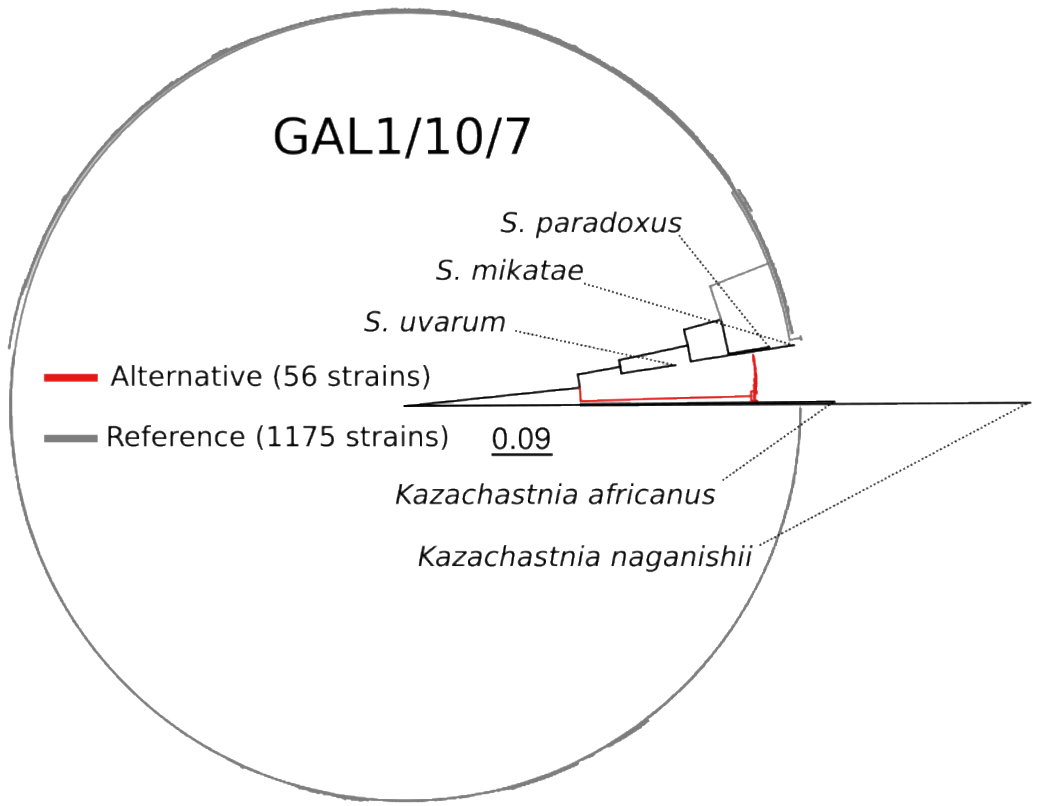


**Supplementary Figure 17: Phylogenetic clustering of the *GAL1/10/7* alleles from the global sample of 1,276 sequenced yeast strains**

Bootstrapped maximum likelihood phylogenetic trees of the *GAL1/10/7* alleles. Gene sequences were successfully extracted from the 1,231 strains with high-quality assemblies in this region. Gene sequences were also extracted from members of the *Saccharyomyces sensu stricto* (*S. uvarum, S. mitakae, S. paradoxus*), and the outgroup species (*Kazachstania africanus* and *Kazachstania naganashii*). Branches are colored according to the genotype of each strain at *GAL1/10/7*; alternative alleles (red), reference alleles (grey), and other species (black). Scale bar shows the estimated number of substitutions per site.

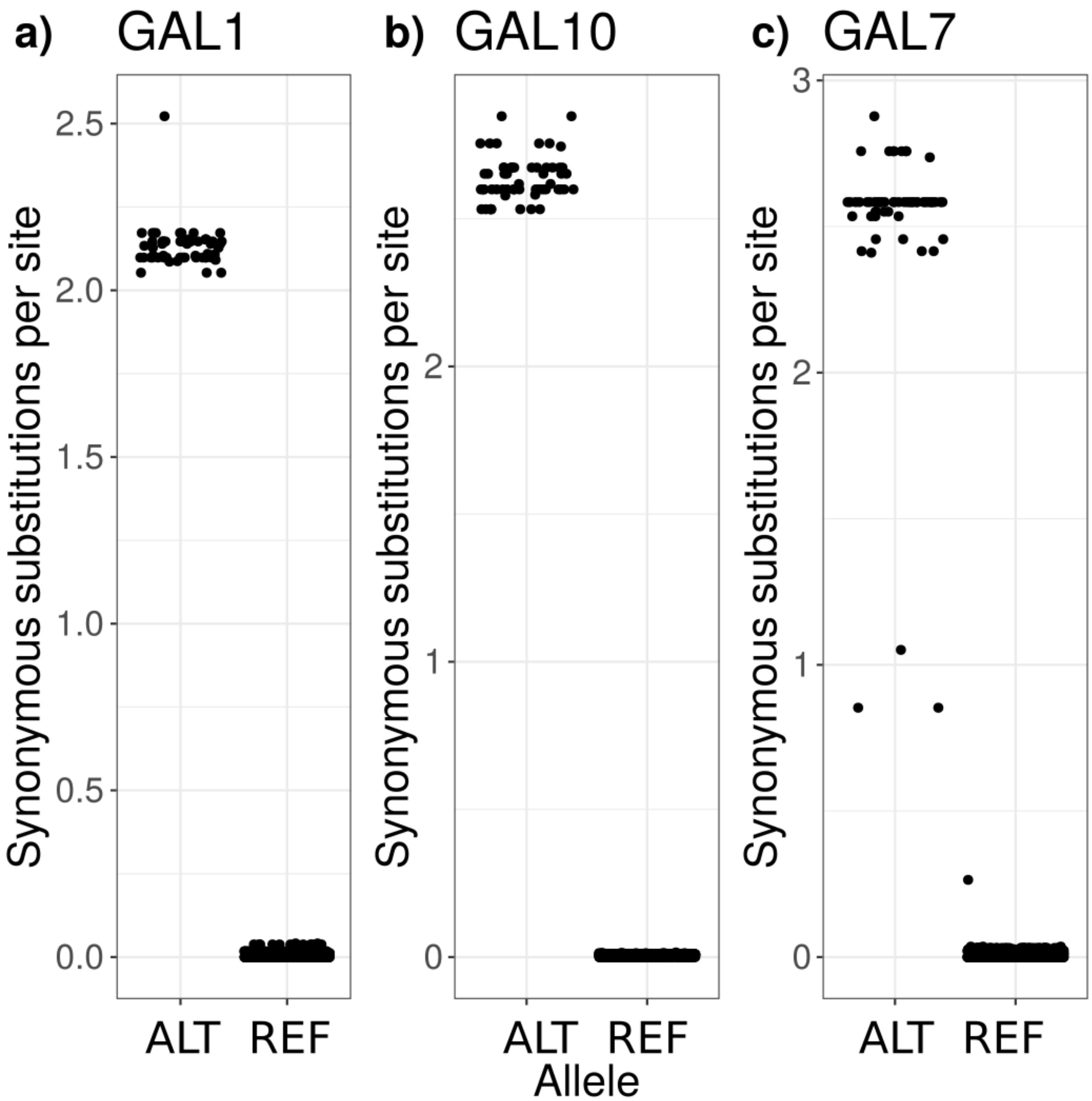


**Supplementary Figure 18: Synonymous substitutions per site (dS) between the genes of SacCer3 and 1,276 sequenced yeast strains for *GAL1*, *GAL10*, and *GAL7*.**

Boxplots of the dS of *GAL1*, *GAL10*, and *GAL7* extracted where possible from the genomes of the 1,276 yeast strains when aligned to the reference genes. Strains are partitioned on the x-axis based on whether they were assigned as having the alternative (ALT) or reference (REF) allele for each gene.

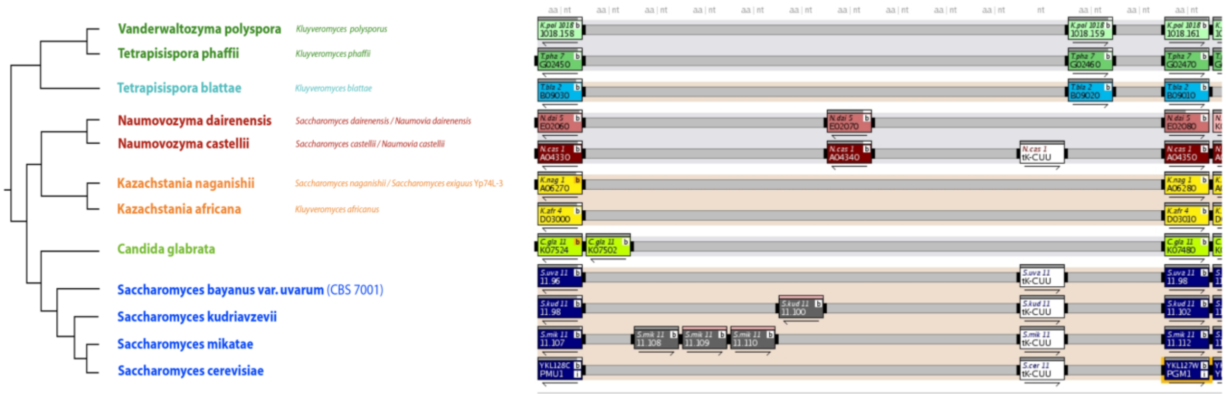


**Supplementary Figure 19: The missing lysine tRNA in the alternative PGM1 promoter is conserved within the *Saccharyomyces sensu stricto*.**

Screenshot of the PGM1 promoter from the yeast gene order browser (*42*). Species from the *Saccharyomyces sensu stricto* (*S. uvarum*, *S. kudriavzevii*, *S. mikatae*, and *S. cerevisiae*) all have a lysine tRNA (tk-CUU) in their *PGM1* promoters whereas the alternative (not shown) *PGM1* promoter does not.

**Table S1: Significant three-way QTL interactions.**

**Table S2: Strains used in this study.**

**Table S3: Primer used in this study.**

**Table S4: Plasmids used in this study.**

**Table S5: Allele-specific expression of a hybrid (CBS2888xYJM981) that is heterozygous for the alternative and reference alleles when grown in 2% glucose medium and transferred to 2% galactose medium.** a) All genes with at least 5 reference and 5 alternate counts, a false-discovery rate adjusted P-value of less than 5% from a binomial test, and a $\log_2$ fold-change of greater than 2 are reported in the table. b) Allele-specific expression for the galactose genes (*GAL1*, *GAL10*, *GAL7*, *GAL2*), and the $\log_2$ fold-change is calculated using the estimated counts of the reference and CBS2888 galactose genes.

**Table S6: Classification of galactose loci in the global collection of 1,276 yeast strains.**

**Table S7: Estimates of the synonymous substitutions per site (dS) between the genes of the reference and CBS2888.**
